# TH1834-Mediated TIP60 Inhibition Protects Against Ischemia/Reperfusion Damage in Human iPSC-Derived Cardiomyocytes and Cardioids

**DOI:** 10.64898/2026.09.17.752307

**Authors:** Zhuocheng Qu, Sakthika Murugesan, Julia H. Ma, Genyu Wang, Nathan R. Tucker, Zhen Ma, Cynthia C. Taub, Xinrui Wang

## Abstract

Ischemia/reperfusion injury limits functional recovery after myocardial infarction through mitochondrial dysfunction, oxidative stress, energetic failure, impaired calcium handling, and contractile dysfunction. We previously identified the acetyltransferase TIP60/KAT5 as a maladaptive regulator of post-ischemic cardiac injury and showed that pharmacologic TIP60 inhibition with TH1834 mitigates myocardial infarction injury in mice. Here, we tested whether TH1834 promotes recovery in complementary 2D and 3D human induced pluripotent stem cell-derived cardiomyocyte (hiPSC-CM) and cardioid (hiPSC-CO) models. Maturation-enhanced hiPSC-CMs subjected to hypoxia/reoxygenation (H/R) reproduced key features of ischemia/reperfusion injury, including mitochondrial loss, oxidative stress, cell death, impaired calcium handling, and contractile dysfunction. RNA sequencing further showed activation of stress programs and suppression of oxidative-metabolic pathways. At a non-arrhythmogenic dose, TH1834 inhibited TIP60 acetyltransferase activity, improved post-H/R survival and beating recovery, preserved mitochondrial and myofibrillar integrity, reduced reactive oxygen species, restored calcium transient kinetics, normalized sarcoplasmic reticulum calcium reserve, and improved contraction and relaxation. A central effect of TH1834 was enhancement of metabolic flexibility: following H/R, treatment restored mitochondrial respiratory capacity while simultaneously increasing glycolytic capacity and reserve. Because CM calcium cycling and mechanical work are highly dependent on energetic supply, this recovery of complementary oxidative and glycolytic capacity provides an energetic basis for improved calcium homeostasis and contractile recovery. Transcriptomic analysis supported this phenotype, showing preferential counter-regulation of H/R-induced stress programs and attenuation of persistent HIF1A-ARNT-PDK1 hypoxic metabolic signaling. In 3D hiPSC-COs, TH1834 similarly enhanced mitochondrial respiratory reserve and glycolytic capacity while improving calcium dynamics, contractile performance, beating recovery, and survival. Together, these findings identify enhanced metabolic flexibility as a central feature of TH1834-mediated protection against H/R, linking restoration of energetic capacity to preservation of calcium signaling and contractile function, supporting TIP60 as a therapeutic target for ischemic heart disease.

## Introduction

Ischemic heart disease remains a leading cause of morbidity and mortality worldwide, and myocardial ischemia/reperfusion injury continues to limit functional recovery after restoration of coronary blood flow. Although reperfusion is essential for salvaging ischemic myocardium, the abrupt return of oxygen and nutrients can paradoxically exacerbate cardiomyocyte (CM) injury through mitochondrial dysfunction, oxidative stress, Ca²⁺ overload, impaired excitation-contraction coupling, sarcomere disruption, and activation of cell death pathways (Hausenloy & Yellon, 2013; Murphy & Steenbergen, 2008). These acute cellular events contribute to persistent contractile dysfunction, adverse remodeling, and progression toward heart failure. Thus, therapeutic strategies that preserve CM survival, mitochondrial-metabolic integrity, Ca²⁺ handling, and contractile recovery after ischemic stress remain a major unmet need.

Human induced pluripotent stem cell–derived cardiac models provide an important platform for studying mechanisms of ischemic injury and therapeutic recovery. Conventional hiPSC-derived CMs (hiPSC-CMs), however, retain fetal-like structural, electrophysiological, and metabolic features that can limit disease-modeling fidelity and may show relative resistance to hypoxic stress (Ewoldt et al., 2025; Feyen et al., 2020). Metabolic maturation strategies that increase oxidative substrate utilization improve hiPSC-CM electrophysiological properties, sarcoplasmic reticulum (SR) Ca²⁺ cycling, sarcomere organization, mitochondrial distribution, and contractile force (Feyen et al., 2020; Li et al., 2025). Importantly, metabolically matured hiPSC-CMs show greater susceptibility to hypoxia-induced mitochondrial dysfunction and cell death, supporting their use as a more clinically relevant human platform for modeling ischemic damage (Peters et al., 2022). Additionally, three-dimensional hiPSC-derived cardioids (hiPSC-COs) provide a complementary platform to 2D hiPSC-CMs by incorporating tissue-like architecture, multicellular organization, extracellular matrix context, diffusion gradients, and integrated contractile behavior. Recent human cardiac organoid studies show that H/R or acute myocardial infarction–like injury recapitulates key ischemic phenotypes, including cell death, oxidative stress, structural remodeling, electrophysiologic dysfunction, and fibrotic responses, supporting their use for therapeutic testing (O’Hern et al., 2025; Song et al., 2024; Zhang et al., 2025).

To model ischemia/reperfusion injury in hiPSC-CMs, hypoxia/reoxygenation (H/R) protocols are widely used to induce changes in beating activity, sarcomere disruption, electrophysiological properties, oxidative stress, metabolic dysfunction, Ca²⁺-handling abnormalities, and impaired recovery (Fernández-Morales et al., 2019; Häkli et al., 2021; Peters et al., 2022). These endpoints are important because ischemic injury is not defined by CM death alone, but by coordinated disruption of the structural, metabolic, Ca²⁺-handling, and contractile systems that sustain beat-to-beat cardiac performance. During ischemia, reduced oxidative phosphorylation limits ATP production, impairing Na⁺/K⁺ ATPase- and SR Ca²⁺ pump SERCA-dependent ion homeostasis and promoting intracellular Na⁺ and Ca²⁺ accumulation; upon reoxygenation, mitochondrial reactive oxygen species (ROS) production, mitochondrial Ca²⁺ loading, and permeability transition further worsen energetic failure and cell injury (Bertero et al., 2024; Chouchani et al., 2014; Murphy & Steenbergen, 2008). Here, the integrated assessment of myofibrillar organization, mitochondrial function, oxidative stress, Ca²⁺ cycling, SR Ca²⁺ reserve, and contraction/relaxation kinetics provides a rigorous functional framework for evaluating cardiac therapeutics.

The lysine acetyltransferase TIP60/KAT5 has emerged as a stress-responsive regulator of cardiac injury and repair. Prior work from our group showed that Tip60 depletion in neonatal CMs reduced DNA damage response signaling, decreased expression of cell-cycle inhibitory programs, increased CM cell-cycle activity, and improved functional recovery after myocardial infarction (Wang et al., 2021). In adult myocardial infarction models, conditional depletion of Tip60 in CMs improved functional recovery, reduced scarring, and protected against post-injury remodeling (Wang et al., 2022). These genetic findings were extended pharmacologically using TH1834, a small-molecule TIP60 inhibitor that mitigated myocardial infarction injury in mice (Wang et al., 2023). More recent work with pentamidine, an FDA-approved antimicrobial agent with TIP60-inhibitory activity from which TH1834 was derived, further supports pharmacologic TIP60 inhibition as a candidate cardioprotective strategy (Wang et al., 2025). However, whether TH1834 directly protects human CMs from H/R injury and improves functional recovery in human 3D cardiac models remains unknown.

In the present study, we evaluated the therapeutic potential of TH1834 in complementary 2D and 3D human iPSC-derived cardiac platforms. We first optimized hiPSC-CM maturation and H/R conditions to establish a model that recapitulates structural, metabolic, oxidative, Ca²⁺-handling, and contractile hallmarks of ischemia/reperfusion injury. We then tested whether TH1834 inhibits TIP60 acetyltransferase activity in mature hiPSC-CMs and whether a non-arrhythmogenic dose improves post-H/R survival and functional recovery. Finally, we extended this approach to hiPSC-COs to determine whether TH1834 preserves Ca²⁺ cycling, contractile performance, mitochondrial function, glycolytic capacity, cell survival, and spontaneous beating recovery in a 3D human cardiac tissue-like context. We hypothesized that pharmacologic inhibition of TIP60 acetyltransferase activity with TH1834 protects human hiPSC-CMs and hiPSC-COs from H/R injury by preserving mitochondrial-metabolic resilience, restoring Ca²⁺ handling, reducing oxidative stress and cell death, and promoting post-ischemic functional recovery.

## Methods

### hiPSC culture and generation of hiPSC-CMs and hiPSC-COs

WTC-derived hiPSC reporter lines, including WTC-GCaMP6f and WTC-mEGFP-ACTN2, were maintained under feeder-free conditions according to Allen Institute for Cell Science protocols. hiPSCs were cultured on growth factor-reduced Matrigel (Corning, 356231)-coated plates (Falcon, 353046) in mTeSR1 medium (STEMCELL Technologies, 85851) at 37°C with 5% CO₂. Cells were passaged using Accutase (Thermo Fisher Scientific/Gibco, A11105-01) and maintained for 24 h after dissociation in mTeSR1 supplemented with ROCK inhibitor Y-27632 (STEMCELL Technologies, 72308).

hiPSC-CMs were generated using small-molecule Wnt modulation adapted from the AICS cardiomyocyte differentiation protocol (Lian et al., 2012). For 2D hiPSC-CM differentiation, hiPSCs were plated on Matrigel-coated 6-well plates on Day −3. Differentiation was initiated on Day 0 using CHIR99021 (Selleckchem, S1263) in RPMI/B27 minus insulin [RPMI/B27(−); Gibco, 11875-093/A18956-01]. After 48 h, medium was changed to IWR1(Sigma-Aldrich, I0161) in RPMI/B27(−). On Day 4, cultures were maintained in RPMI/B27(−), followed by RPMI/B27 with insulin [RPMI/B27(+); Gibco, 11875-093/17504-044] on Day 6. Media were changed every 48 h, and spontaneous beating was monitored beginning around Days 7–11.

For 2D hiPSC-CM purification, cultures were switched on Day 12 to glucose-free RPMI/B27(+) [RPMI(-)/B27(+), Gibco 11879-020/17504-044]. On Day 15, cells were replated in RPMI(-)/B27(+) supplemented with 10 μM Y-27632. On Day 16, hiPSC-CMs were maintained in either control RPMI/B27(+) medium or maturation medium with twice-weekly media changes. Maturation medium was modified from published metabolic maturation protocols (Feyen et al., 2020; Peters et al., 2022; Li et al., 2025) and consisted of glucose-free DMEM (Thermo Fisher Scientific, 11966025) supplemented with 3 mM glucose (Sigma-Aldrich, G8270), 5 mM creatine monohydrate (Thermo Fisher Scientific, 226791000), 2 mM taurine (Sigma-Aldrich, T0625), 2 mM L-carnitine (Thermo Fisher Scientific, 230281000), 1× non-essential amino acids (Thermo Fisher Scientific, 11140-050), 0.5% AlbuMAX (Gibco, 11021-029), 1× B27 (Gibco, 17504-044), and 1% KOSR (Thermo Fisher Scientific). 2D hiPSC-CMs were cultured in maturation medium for at least 6 days before H/R experiments.

For 3D hiPSC-CO generation, dissociated hiPSCs were seeded into Costar ultra-low attachment 96-well plates (Costar, 7007) in mTeSR1 supplemented with Y-27632 and centrifuged at 300 × g for 5 min to promote aggregation. Cardiac differentiation was initiated using CHIR99021 followed by IWR1, as described above and adapted from our previous protocol (Hoang et al., 2018; Song et al., 2025). hiPSC-COs were maintained with media changes every 48 h and monitored for spontaneous beating. hiPSC-COs were changed to maturation medium 72 h before hypoxia (or normoxia control) exposure and studied no earlier than Day 30 post-differentiation.

### Hypoxia and reoxygenation

H/R injury was induced using a hypoxia incubator (Biospherix, ProOX C21) with continuous oxygen monitoring. Maturation medium containing either 0.5 mM or 3 mM glucose was equilibrated in the 0.5% O₂ chamber for at least 24 h before use. hiPSC-CMs or hiPSC-COs were exposed to 0.5% O₂ at 37°C with 5% CO₂ for 24 h. After hypoxia, cultures were reoxygenated at 21% O₂, 37°C, and 5% CO₂ for 30 min unless otherwise indicated. Samples were then processed immediately for designated assays.

### TH1834 treatment

TH1834 (Sigma-Aldrich SML3018) was dissolved in PBS to prepare a 1 mg/mL stock solution and stored in aliquots at −20°C protected from light. Working solutions were freshly prepared by diluting TH1834 stock into the appropriate culture medium immediately before treatment. Vehicle control cultures received an equivalent volume of PBS. For dose-response studies, hiPSC-CMs were treated with increasing concentrations of TH1834 under normoxic conditions, and spontaneous beating activity was assessed to identify a non-arrhythmogenic working concentration. For H/R experiments, hiPSC-CMs or hiPSC-COs were treated with 5 µM TH1834 or vehicle during the 24 h hypoxia period and maintained in the same treatment condition during reoxygenation.

### Caspase-3/7, MitoTracker, and CellROX fluorescence assays

Caspase-3/7 activity, functional mitochondria, and ROS were measured in hiPSC-CMs and hiPSC-COs using fluorescence-based live-cell assays adapted from the manufacturer’s protocols. CellEvent Caspase-3/7 Red Detection Reagent (Invitrogen, C10430) was prepared fresh on the day of use. The reagent was reconstituted in PBS to generate a 100x stock solution, diluted in culture medium to 1x working solution immediately before use. CMs or COs were incubated for 40 min or 2h at 37°C before fluorescence measurement without washing using excitation/emission settings of 590/610 nm. Functional mitochondria were detected using MitoTracker Orange CMTMRos (Invitrogen, M7510). Lyophilized MitoTracker Orange CMTMRos was dissolved in anhydrous DMSO to prepare a 1 mM stock solution and diluted in prewarmed culture medium to 100nM working solution immediately before use. CMs or COs were incubated with MitoTracker staining solution for 40 min or 2h at 37°C, washed with prewarmed medium, and fluorescence was measured using excitation/emission settings of 554/576 nm. ROS were measured using CellROX Deep Red Reagent (Invitrogen, C10422). CellROX reagent was diluted in culture medium to 5 µM final concentration immediately before use. CMs or COs were incubated for 30 min or 2h at 37°C, washed with PBS, fluorescence was measured using excitation/emission settings of 640/665 nm. Fluorescence readings for all assays were obtained using a FlexStation 3 Benchtop Multi-Mode Microplate Reader (Molecular Devices) and normalized to DNA (DAPI) or mitochondria (MitoTracker) content as indicated.

### Immunostaining of hiPSC-CMs and hiPSC-CO frozen sections

For 2D hiPSC-CM immunostaining, dissociated CMs were replated onto Matrigel-coated 12-mm round glass coverslips (Electron Microscopy Sciences, 72231-01) or Matrigel-coated 22 mm × 22 mm square glass coverslips (Alkali Scientific, SM2222). Following cell attachment and spreading, cultures were washed with PBS and fixed in 4% paraformaldehyde (PFA) for 15 min at room temperature. After fixation, cells were washed with PBS and either stained immediately or stored in PBS at 4°C until further processing. For hiPSC-CO frozen sections, COs were washed twice with PBS and fixed in 4% PFA for 30 min at room temperature. After fixation, samples were washed twice with PBS. Fixed hiPSC-COs were embedded in Tissue-Tek O.C.T. Compound (Sakura, 4583) using Tissue-Tek cryomolds (Sakura, 4565). The embedded samples were initially cooled in a cryostat pre-cooled to −20°C for approximately 10 min, followed by an additional 10 min on the quick-freeze shelf under a pre-cooled heat extractor to ensure complete freezing throughout the OCT block before cryosectioning. Frozen blocks were then mounted onto specimen discs using O.C.T. Compound and cryosectioned at a thickness of 10 µm.

CMs or CO sections were permeabilized with 0.5% Triton X-100 in PBS for 10 min at 4°C, washed three times with PBS for 5 min each, and blocked with 3% bovine serum albumin (BSA) in PBS for 1h at room temperature. Samples were incubated with primary antibodies diluted in blocking buffer overnight at 4°C. After primary antibody incubation, samples were washed three times with PBS for 5 min each, incubated with species-appropriate Alexa Fluor 488-, 555-, or 647-conjugated secondary antibodies (Invitrogen A11029/A11034, A21429/A21424/A78945, A21245/A21236; 1:400) for 1 h at room temperature, washed three times with PBS for 5 min each, counterstained with DAPI diluted 1:2000 for 10 min at room temperature, washed once with PBS, and mounted using ProLong Diamond Antifade Mountant (Invitrogen, P36970). Imaging was performed on a Leica SP8 confocal microscope (Upstate/Leica Center of Excellence) and image analysis using LAS X and ImageJ (Fiji) v2.14.0 software. Primary antibodies used were: rat anti-α-tubulin (Invitrogen/Thermo Fisher Scientific, MA1-80017; 1:500), mouse anti-TOMM20 (Abcam, ab56783; 1:2000), rabbit anti-acetylated histone H2A.Z (Cell Signaling Technology, 75336S; 1:400), rabbit anti-acetylated histone H4 (Abcam, ab177790; 1:5000), rabbit anti-MLC-2v (Proteintech, 10906-1-AP; 1:400), rat anti-Ki67 (Invitrogen 14-5698-82; 1:250), and rabbit anti-α-actinin (Abcam, ab68167; 1:500).

### Ca^2+^ transient and contractility measurements

Intracellular Ca²⁺ transients and contractile motion were measured in GCaMP-expressing hiPSC-CMs and hiPSC-COs using an IonOptix calcium and contractility acquisition system as previously described (Wang and Fitts, 2018). hiPSC-CMs were washed and maintained in Tyrode’s buffer (1.8mM CaCl_2_, 130 mM NaCl, 4 mM KCl, 0.33 mM NaH₂PO₄, 1 mM MgCl₂, 10 mM HEPES, 10 mM glucose, 20 mM taurine, pH 7.4) at 37°C and field-stimulated at the indicated pacing frequency. For β-adrenergic stimulation experiments, hiPSC-CMs were paced at 2 Hz and exposed to 1 µM isoproterenol (Sigma-Aldrich, PHR2722) during acquisition. To assess SR Ca²⁺ store, a 20mM caffeine bolus was applied to cells that were responsive to 1Hz stimulation to induce total SR Ca²⁺ release. hiPSC-COs were maintained in culture medium at 37°C and recorded under spontaneous beating conditions without electrical stimulation.

GCaMP fluorescence was recorded using GFP-compatible excitation and emission settings. Contractile motion of hiPSC-CMs was acquired simultaneously using the CytoMotion module. Contractility was quantified from brightfield image sequences based on pixel intensity and pixel correlation changes within a selected region of interest. For hiPSC-CMs, only contracting region within single cell was selected; for hiPSC-COs, the region of interest was positioned over an area with a clear center of contraction. Background-corrected Ca²⁺ and contraction traces were analyzed using CytoSolver software. Ca²⁺ transient parameters included resting fluorescence, peak amplitude, maximal departure/rise rate, maximal decay/return rate, time to peak, and time to 50% and 90% return to baseline. Contractility parameters included contraction amplitude, maximal contraction velocity, maximal relaxation velocity, and time to 50% and 90% return to baseline. Stable consecutive beats were averaged for each cell/field and normalized to the corresponding baseline. Ca²⁺ and contraction data were collected from at least three individual batches and were normalized to the averaged values of the normoxia group within each experimental batch.

### Seahorse Metabolic Assay

Mitochondrial respiration and glycolytic function were evaluated in 2D hiPSC-CMs and hiPSC-COs by measuring oxygen consumption rate (OCR, pmol/min) and extracellular acidification rate (ECAR, mpH/min), respectively, using an XF96 Extracellular Flux Analyzer (Agilent). The day before the assay, 200 µL of XF Calibrant was added to each well of the XF96 utility plate, and the sensor cartridge was hydrated overnight in a 37°C non-CO₂ incubator. For 2D hiPSC-CMs, cells were seeded onto Matrigel-coated XF96 cell culture microplates at the designated density and cultured until the time of assay. For hiPSC-COs, individual COs were transferred into Matrigel-coated XF96 assay plate wells and centered within each well before measurement. One hour before the assay, culture medium was replaced with Seahorse assay medium containing unbuffered RPMI supplemented with the designated drug treatment. For the mitochondrial stress test, assay medium was supplemented with 1 mM pyruvate, 2 mM L-glutamine, and 10 mM glucose. For the glycolytic stress test, assay medium was supplemented with 2 mM L-glutamine. Plates were incubated for 1 h at 37°C in a non-CO₂ incubator before analysis.

For the mitochondrial stress test, OCR was measured at baseline and after sequential automated injections of oligomycin, carbonyl cyanide-p-trifluoromethoxyphenylhydrazone (FCCP), and rotenone + antimycin A, yielding final concentrations of 2 µM oligomycin, 2 µM FCCP, and 1 µM rotenone + antimycin A. OCR parameters were calculated as follows: basal respiration was defined as the last rate measurement before oligomycin injection minus non-mitochondrial respiration; maximal respiration was defined as the maximum rate measurement after FCCP injection minus non-mitochondrial respiration; proton leak was defined as the minimum rate measurement after oligomycin injection minus non-mitochondrial respiration; and non-mitochondrial respiration was defined as the minimum rate measurement after antimycin A injection. For the glycolytic stress test, ECAR was measured at baseline and after sequential automated injections of glucose, oligomycin, and 2-deoxy-D-glucose (2-DG), yielding final concentrations of 20 mM glucose, 2 µM oligomycin, and 50 mM 2-DG. ECAR parameters were calculated as follows: glycolysis was defined as the maximum rate measurement before oligomycin injection minus the last rate measurement before glucose injection; glycolytic capacity was defined as the maximum rate measurement after oligomycin injection minus the last rate measurement before glucose injection; non-glycolytic acidification was defined as the last rate measurement before glucose injection; and glycolytic reserve was calculated as glycolytic capacity minus glycolysis. For both 2D hiPSC-CMs and hiPSC-COs, OCR and ECAR values were normalized to DNA content (DAPI) measured from the corresponding well.

### RNA isolation and Sequencing

Maturation-enhanced hiPSC-CMs were studied under three conditions: normoxia, hypoxia/reoxygenation (H/R), and H/R with TH1834 treatment (H/R + TH). For H/R, cells were exposed to 0.5% O₂ for 24 h in maturation medium containing 3 mM glucose followed by 30 min reoxygenation at 21% O₂. TH1834 (5 µM) was present during the 24-h hypoxic exposure and reoxygenation period. Two independent biological replicates were analyzed for each condition (n=2 per group; six samples total). RNA was collected at the end of the 30-min reoxygenation period; normoxic samples were collected at the corresponding culture time point.

Total RNA was isolated using RNA mini (Invitrogen 12183018A) isolation kit according to the manufacturer’s instructions. RNA concentration and purity were assessed using NanoDrop, and RNA integrity was evaluated using Agilent TapeStation. rRNA-depleted strand-specific sequencing libraries were generated using the NEBNext Ultra II Directional RNA Library Prep Kit for Illumina (New England Biolabs, E7760). Libraries were sequenced on an Illumina NextSeq 2000 using single-end 100-bp reads to an average depth of 55 million reads per sample.

Sequencing reads were assessed using FastQC v0.12.1 and processed for adapter and low-quality sequence removal using fastp v1.3.3. Quality-control metrics were summarized using MultiQC v1.35. Reads were aligned to the GENCODE GRCh38.p14 primary genome assembly using STAR v2.7.11b with the GENCODE release 50 comprehensive gene annotation. Reverse-stranded gene-level raw counts were generated directly using STAR --quantMode GeneCounts. Transcript abundance was additionally expressed as transcripts per million (TPM) and fragments per kilobase per million (FPKM) using GENCODE release 50 exon-union gene lengths. TPM and FPKM values were used only for descriptive visualization of gene-expression levels; all statistical differential-expression analyses were performed using integer raw gene counts.

### Differential Gene-expression Analysis

Differential expression was analyzed from raw reverse-stranded gene counts using DESeq2 v1.42.1 (Love et al., 2014). Analysis was restricted to the six biological samples comprising normoxia, H/R, and H/R+TH1834 groups (n=2/group). Genes were retained when raw counts were ≥10 in both biological replicates of at least one of the three experimental conditions. A single DESeq2 model with experimental condition as the design factor (design = ∼ group) was fitted across all samples, providing shared normalization and dispersion estimation for the three experimental groups. Three prespecified contrasts were evaluated: H/R versus normoxia, H/R+TH1834 versus H/R, and H/R+TH1834 versus normoxia.

Statistical significance was determined using DESeq2 Wald tests followed by Benjamini– Hochberg correction for multiple comparisons. Independent filtering was enabled separately for each contrast using alpha = 0.05. Genes with an adjusted P value (padj) <0.05 were considered statistically significant. Log₂ fold-change estimates were subjected to DESeq2 shrinkage using the normal prior. The DESeq2 model included all annotated gene biotypes that passed the expression filter, whereas downstream manuscript DEG visualization, rescue analysis, and pathway analysis were restricted to genes annotated as protein-coding in GENCODE release 50. For characterization of the magnitude of the H/R injury response, protein-coding genes with padj <0.05 and an absolute shrunken log₂ fold change ≥1 (≥2-fold change) were classified as high-magnitude H/R differentially expressed genes (DEGs). This gene set was used for focused H/R DEG visualization and directional over-representation analyses. For analyses of the broader H/R-responsive transcriptome and treatment-mediated rescue, protein-coding H/R-responsive genes with padj <0.05 were retained without an additional fold-change cutoff.

Variance-stabilizing transformation (VST) of raw counts was performed using DESeq2 with blind = FALSE and the same unified six-sample model. VST-transformed values were used for principal-component analysis, sample-level visualization, heatmaps, and calculation of group-level expression trajectories. TPM and FPKM values were used only for descriptive reporting of transcript abundance and were not used for statistical testing.

### Transcriptomic Rescue Analysis

To determine whether TH1834 counter-regulated H/R-induced transcriptional changes, rescue analysis was performed using genes significantly altered by H/R relative to normoxia (DESeq2 padj <0.05), without an additional fold-change threshold. For each gene, the H/R effect was defined by the H/R-versus-normoxia contrast, and the treatment effect was defined by the H/R + TH1834-versus-H/R contrast. Normal-prior shrunken log₂ fold changes were used to determine the directions of these effects, and mean VST-transformed expression was calculated for the normoxia, H/R, and H/R + TH1834 groups.

A gene was classified as showing directional rescue when three conditions were satisfied: (1) the normal-prior shrunken log₂ fold changes for H/R + TH1834 versus H/R and H/R versus normoxia were opposite in direction; (2) the corresponding group-mean VST expression changes were also in opposite directions, indicating movement toward the normoxic level after TH1834 treatment; and (3) the absolute difference between the mean expression levels of H/R + TH1834 and normoxia was smaller than that between untreated H/R and normoxia. Strict rescue genes additionally required padj <0.05 for both the H/R versus normoxia and H/R+TH1834 versus H/R contrasts.

The magnitude of rescue was quantified from VST-transformed group means as: Rescue fraction = (mean H/R+TH1834 – mean H/R) / (mean Normoxia – mean H/R). A rescue fraction of 0 indicates no movement toward normoxia, whereas 1(100%) represents movement to the normoxic mean. Directionally rescued genes were descriptively classified as minimal rescue (>0–20%), partial rescue (20–80%), near-normoxic trajectory (80–105%), or overshoot beyond the normoxic mean (>105%). These categories describe the magnitude of the expression trajectories and do not represent formal equivalence tests. Accordingly, a nonsignificant H/R+TH1834 versus normoxia comparison was described as “no statistically detectable difference” and was not interpreted as proof of equivalent expression.

### Statistical Analysis

Two hiPSC lines were used and the individual points depicted in scatter plots represent the number of independent experiments performed. Results are shown as mean ± SEM. Statistical analysis was performed using Prism 10 (GraphPad) software. Normality was assessed using a Shapiro-Wilk test. If data comprised two normally distributed groups, a parametric unpaired two-tailed Student’s *t* test was performed to determine significant differences. If data comprised two non-normally distributed groups, a nonparametric two-tailed Wilcoxon/Mann-Whitney U test was performed. For three or more groups and the assessment of one parameter, one-way ANOVA statistical test was used. Dunnett’s multiple comparison test was used as post hoc analysis to determine significance.

## Results

### Maturation enhances hiPSC-CM structural organization and sensitivity to ischemic stress

To establish a human CM platform suitable for modeling ischemia/reperfusion-like injury, we first developed a maturation media modified from previously reported metabolic and functional maturation strategies, including low-glucose, fatty acid–enriched, oxidative-substrate–supporting, and Ca²⁺-optimized approaches (Feyen et al., 2020; Peters et al., 2022; Li et al., 2025). These studies collectively demonstrate that metabolic substrate switching, fatty acid supplementation, increased Ca²⁺ availability, micropatterning/nanopatterning, and electro-mechanical maturation cues can improve hiPSC-CM structural, metabolic, electrophysiological, and contractile phenotypes. In our system, mature CM structural and contractile phenotypes were evident as early as day 3 after application of maturation media, as shown in Supplemental Videos 1-2. By day 6, CMs from maturation media exhibited well-aligned sarcomeres and robust shortening (Supplemental Videos 3-4). Compared with control RPMI/B27(+) media, maturation media improved cell alignment, granularity, sarcomere length, and contractile behavior on days 6, 9, and 12, and sustained contractile function was maintained through day 60 post-differentiation (Supplemental Videos 5-8). Consistent with these live-cell observations, maturation enhanced hiPSC-CMs displayed phenotypic features of enhanced structural and metabolic maturation, including tighter sarcomere alignment marked by sarcomeric α-actinin (ACTN2), robust expression of the ventricular isoform of myosin light chain (MLC-2v), and redistribution of functional mitochondria (MitoTracker) toward myofibrillar networks rather than the predominantly perinuclear mitochondrial clustering observed in immature cells (Fig. 1A).

**Figure 1.**
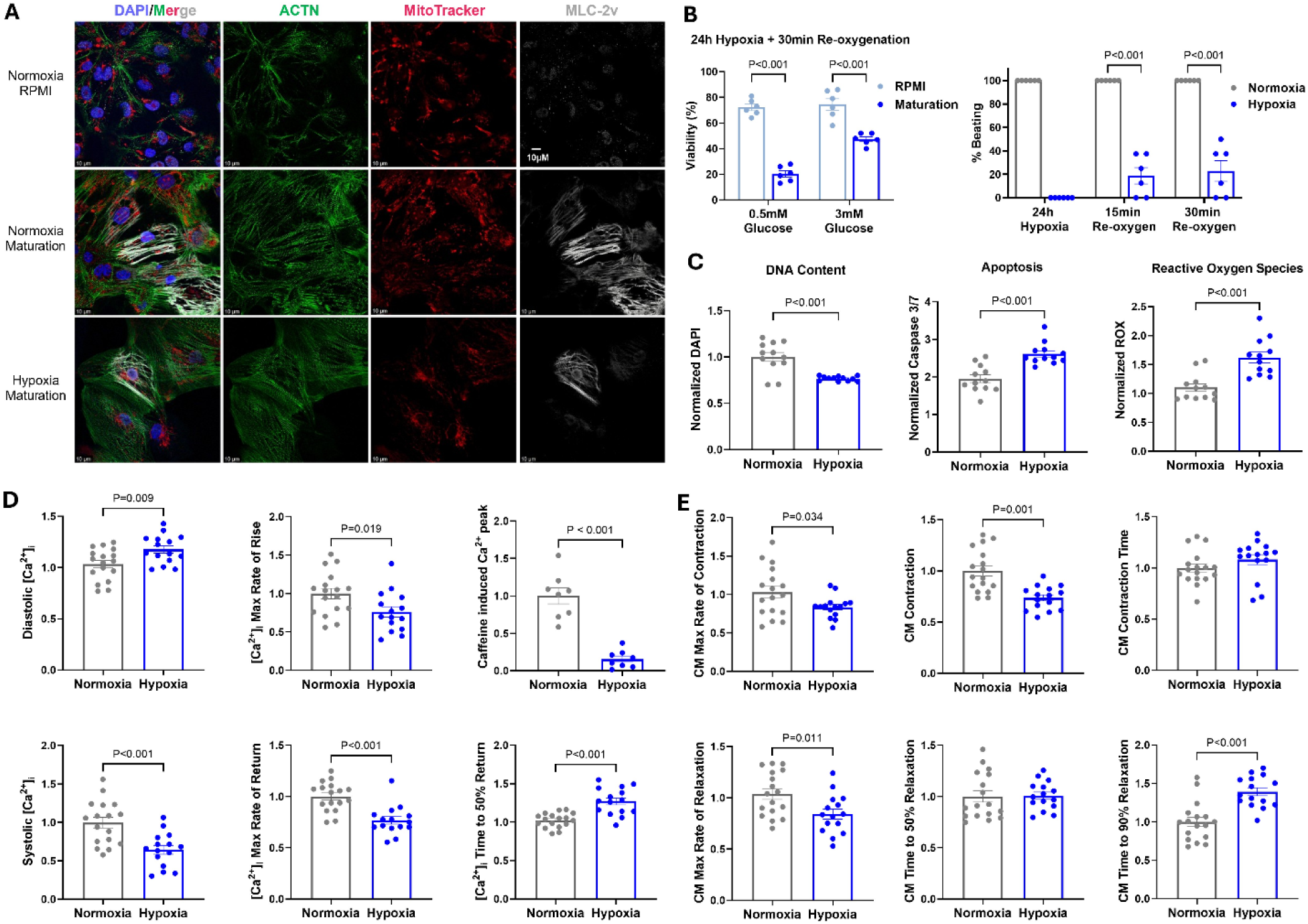
Maturation-enhanced hiPSC-CMs establish a hypoxia/reoxygenation model that recapitulates ischemia/reperfusion injury. **A.** Representative immunofluorescence images of hiPSC-CMs cultured in standard RPMI/B27(+) medium or maturation medium under normoxic conditions, and hiPSC-CMs cultured in maturation medium after H/R. Cells were stained for DAPI, sarcomeric α-actinin (ACTN), MitoTracker, and ventricular myosin light chain 2 (MLC-2v). Compared with RPMI/B27(+) medium, maturation medium enhanced sarcomere alignment, increased MLC-2v expression, and promoted redistribution of functional mitochondria toward myofibrillar networks. H/R disrupted myofibrillar organization and reduced MitoTracker signal in maturation-enhanced hiPSC-CMs. Scale bars, 10 µm. **B.** Optimization of H/R injury severity in hiPSC-CMs. After 24 h hypoxia at 0.5% O₂ followed by 30 min reoxygenation, maturation-enhanced hiPSC-CMs showed glucose-dependent injury severity. Hypoxia in 0.5 mM glucose caused severe loss of viability, whereas 3 mM glucose produced an intermediate injury phenotype. Spontaneous beating was arrested after 24 h hypoxia and partially recovered after 15 and 30 min reoxygenation. **C.** H/R recapitulated core cellular hallmarks of ischemia/reperfusion injury in maturation-enhanced hiPSC-CMs, including reduced DNA content/DAPI signal, increased caspase-3/7 activation, and elevated reactive oxygen species. **D.** H/R impaired intracellular Ca²⁺ handling in hiPSC-CMs paced at 1 Hz. H/R increased resting diastolic [Ca²⁺]i, reduced systolic [Ca²⁺]i peak, decreased maximal [Ca²⁺]i rise and return rates, and prolonged time to 50% return to baseline. H/R also decreased caffeine-induced sarcoplasmic reticulum (SR) Ca²⁺ release. **E.** H/R impaired single-cell contractile performance in hiPSC-CMs paced at 1 Hz. H/R depressed contraction, reduced maximal rates of contraction and relaxation, and prolonged time to 90% relaxation. Data are presented as mean ± SEM. Ca²⁺ transient and contraction data were normalized to the averaged values of the normoxia group within each experimental batch. Statistical comparisons are indicated in each panel. RPMI, RPMI/B27(+) medium; Maturation, maturation medium; H/R, 24 h hypoxia at 0.5% O₂ with 3 mM glucose followed by 30 min reoxygenation at 21% O₂.

Total RNA sequencing of normoxic maturation-enhanced hiPSC-CMs provided complementary molecular evidence of a ventricular, maturation-associated phenotype. Cells robustly expressed sarcomeric and ventricular genes (Supplemental table 1), including MYH7, MYBPC3, MYL2, ACTN2, IRX4, and HEY2, together with genes supporting Ca²⁺ handling and electrical coupling, including ATP2A2, RyR2, PLN, GJA1, JPH2, SLC8A1, SCN5A, and KCNJ2. The energetic program included high expression of CKM and FABP3, with detectable CKMT2. This coordinated expression across sarcomeric, Ca²⁺-handling, electrophysiological, and metabolic modules resembles maturation-associated transcriptional programs reported in metabolically and physiologically matured hiPSC-CMs (Feyen et al., 2020; Zhang et al., 2026). However, developmental isoforms remained prominent, including MYH6 and TNNI1 relative to MYH7 and TNNI3, respectively, indicating that the cells retain features of incomplete developmental maturation. Thus, the transcriptomic profile supports a maturation-enhanced ventricular phenotype rather than complete acquisition of an adult ventricular transcriptional state.

We next evaluated H/R conditions that would produce reproducible injury while preserving a dynamic range for detecting therapeutic rescue. When exposed to 24h hypoxia at 0.5% O₂ in low-glucose medium, hiPSC-CMs from maturation media showed glucose-dependent injury severity. Hypoxia in 0.5 mM glucose caused severe loss of viability, whereas 3 mM glucose produced an intermediate injury phenotype with measurable residual viability and recovery potential for spontaneous beating after 30 min re-oxygenation at 21% O₂ (Fig. 1B). Following 24h hypoxia and 30min re-oxygenation with 3 mM glucose, CMs from maturation media exhibited disrupted myofibrillar organization and reduced mitochondrial content, indicating vulnerability to ischemic stress (Fig. 1A). In contrast, CMs from standard RPMI/B27(+) media were comparatively resistant to the same ischemic stress conditions, supporting the use of maturation enhanced hiPSC-CMs for modeling clinically relevant post-ischemic injury.

### H/R recapitulates hallmarks of ischemia/reperfusion injury in hiPSC-CMs

Based on the optimization studies, we selected 24h hypoxia at 0.5% O₂ with 3 mM glucose followed by 30min reoxygenation at 21% O₂ as the H/R condition for subsequent experiments, with hypoxia initiated at least 6 days after switching to maturation media. Besides disrupted sarcomere structure, decreased number of functional mitochondria, and spontaneous beating arrest, this protocol recapitulated addition key features of ischemia/reperfusion injury, including reduced cell number as measured by viability and DNA content, increased apoptosis as reflected by caspase-3/7 activation, and elevated oxidative stress as indicated by increased ROS production (Fig. 1A-C).

We further assessed whether this H/R condition reproduced functional hallmarks of ischemia/reperfusion injury at single cell level. H/R markedly impaired intracellular calcium ([Ca²⁺]_i_) handling in hiPSC-CMs stimulated at 1Hz, as shown by elevated resting diastolic [Ca²⁺]_i_, lower systolic [Ca²⁺]_i_ peak, reduced [Ca²⁺]_i_ transient rise and decay rates, and delayed return of [Ca²⁺]_i_ to 50% of baseline. H/R also decreased SR Ca²⁺ store reflected by caffeine-induced Ca²⁺ release (Fig. 1D). These findings indicate that H/R disrupts [Ca²⁺]_i_ homeostasis, elevates diastolic [Ca²⁺]_i_ load, slows [Ca²⁺]_i_ cycling kinetics, and delays clearance during post-ischemic recovery. Consistent with impaired [Ca²⁺]_i_ handling, H/R also impaired single cell contractile performance in hiPSC-CMs paced at 1Hz. H/R compromised contraction, diminished rate of relaxation, and prolonged relaxation time, indicating depressed contractile force generation and impaired cross-bridge cycling dynamics after ischemic stress (Fig. 1E). Together, these data establish a hiPSC-CM H/R injury model that captures structural disruption, mitochondrial loss, oxidative stress, cell death, [Ca²⁺]_i_ handling dysfunction, and contractile impairment while maintaining responsiveness to electrical pacing and sufficient dynamic range to evaluate cardioprotective interventions such as TH1834 treatment.

### H/R induces a broad cellular stress response while suppressing CM contractile and metabolic programs

To further define the molecular response of maturation-enhanced hiPSC-CMs to H/R injury, we performed total RNA sequencing following the optimized H/R protocol. H/R altered 3,918 genes more than 2-fold relative to normoxia, including 1,464 upregulated and 2,454 downregulated genes (Fig. 2A-C). Among the most strongly induced protein-coding genes were *HSPA6, MT1H, MT1M, MT1G, MT2A, HSPA1B, HMOX1, MT1E, and MT1X*, revealing a prominent metallothionein, heat-shock, and oxidative-stress response. H/R also increased established hypoxia-responsive and metabolic-adaptation genes, including *NDRG1, PFKFB4, ADM, DNAJB1, and EGLN1* (Fig. 2A). In parallel with activation of stress-adaptation genes, H/R markedly suppressed transcripts associated with CM structural and functional identity. These included *LRRC10, IRX4, MYL2, MYL7, MYH7, ATP2A2*, and *SELENON*. Genes supporting oxidative and fatty-acid metabolism were also reduced, including *HMGCS2, PPARA, and APOA1* (Fig. 2A, B). Thus, the H/R transcriptome was characterized not only by induction of an adaptive hypoxia/stress response but also by broad suppression of transcriptional programs that support cellular metabolism and cardiac function. This pattern is consistent with transcriptomic studies of infarcted mouse and human myocardium showing stress-responsive CM states accompanied by regional remodeling of metabolic and cardiac functional programs (Calcagno et al., 2022; Yamada et al., 2022; Kuppe et al., 2022).

**Figure 2.**
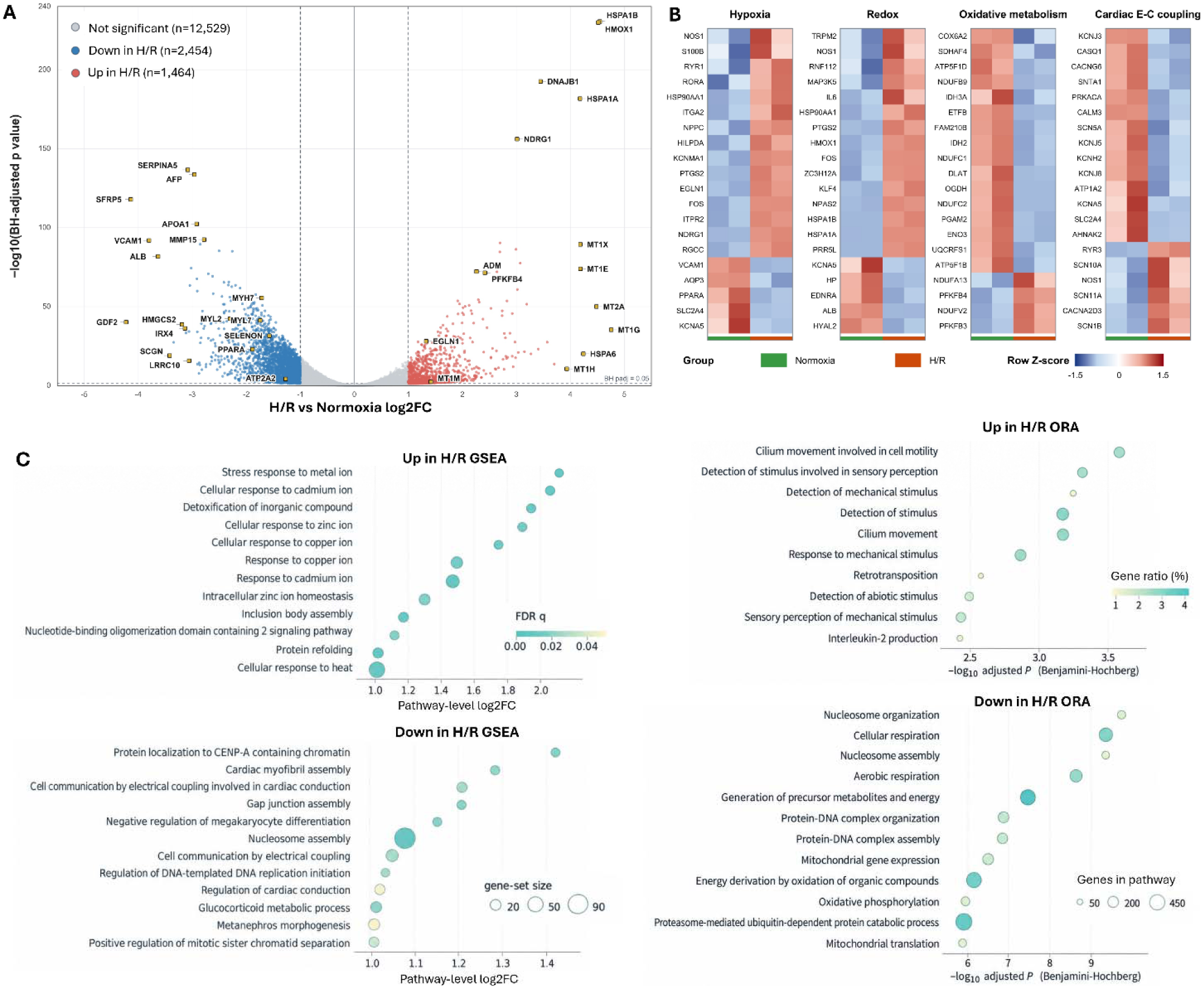
Hypoxia/reoxygenation induces a broad cellular stress response while suppressing CM contractile and metabolic transcriptional programs. **A.** Volcano plot of differential gene expression between H/R and normoxia. The x-axis shows DESeq2 shrunken log2 fold change and the y-axis shows −log10 Benjamini–Hochberg-adjusted P value. DEGs increased or decreased by H/R with adjusted P<0.05 and |shrunken log2FC|≥1 are shown in red and blue, respectively; gray denotes genes not meeting both thresholds. Gold squares indicate selected genes highlighted for biological relevance. **B.** Heatmaps of H/R-associated changes across four functional gene modules: hypoxia response, redox regulation, oxidative metabolism, and cardiac excitation–contraction coupling. Module genes were defined using human GO annotations and intersected with protein-coding H/R DEGs. Genes with adjusted P<0.05 and |shrunken log2FC|≥1 were retained, and twenty genes per module were prioritized according to the magnitude and direction of the H/R-associated expression change. Heatmap values represent row-wise Z-scores of variance-stabilizing transformation (VST)-normalized expression across samples. **C.** Pathway enrichment analysis of the H/R transcriptional response. Over-representation analysis (ORA) was performed separately on H/R-upregulated and H/R-downregulated DEGs using GO Biological Process terms restricted to 15–500 genes; significant terms were defined by BH-adjusted P<0.05, and the 12 most significant terms in each direction are shown. Ranked gene set enrichment analysis (GSEA) was performed using all 16,447 tested genes ranked by the DESeq2 Wald statistic. Gene sets were restricted to 15–500 genes and considered significant at FDR q≤0.05 with pathway-level |mean log2FC|>1. Under these criteria, 24 gene sets met significance, with all 12 positively and 12 negatively enriched pathways shown.

Pathway analyses further demonstrated this coordinated response (Fig. 2B, C). Over-representation analysis (ORA) of H/R-upregulated genes identified enrichment of stimulus- and mechanical-responses and ciliary processes, whereas H/R-downregulated genes were strongly enriched for cellular and aerobic respiration, oxidative phosphorylation, mitochondrial gene expression/translation, and nucleosome/protein–DNA organization. Complementary ranked GSEA resolved more specific directional changes with increased metal-ion and proteotoxic stress responses and decreased cardiac myofibril assembly, gap-junction/electrical coupling, and cardiac conduction pathways, providing molecular support for the structural and excitation– contraction abnormalities observed functionally. Together, these analyses define a dual H/R transcriptional phenotype comprising activation of cellular stress responses together with suppression of metabolic, myofibrillar, and electrical-coupling programs. These molecular changes closely parallel the H/R-induced loss of functional mitochondria and myofibrillar organization, increased oxidative stress, impaired SR Ca²⁺ reserve and Ca²⁺ transient kinetics, and depressed contraction/relaxation observed in Figure 1A-E.

### TH1834 inhibits TIP60 acetyltransferase activity and promotes hiPSC-CM survival after H/R

We next tested whether the TIP60 inhibitor TH1834 engages its intended target in hiPSC-CMs and whether it improves post-ischemic recovery. Immunostaining showed that overnight TH1834 treatment at 5 µM reduced acetylation of the TIP60 target histones H4 (H4ac^K5/8/12/16^) and variant H2A.Z (H2A.Zac^K4/7^), supporting on-target suppression of TIP60 lysine acetyltransferase activity in the setting of preserved sarcomeric structure and mitochondrial labeling (Fig. 3A). We then assessed the dose-dependent effects of TH1834 on spontaneous beating behavior under basal conditions. TH1834 at 5 µM or lower had little to no effect on beating rate, whereas 10 µM produced arrhythmogenic effects, indicating that higher-dose treatment adversely affects electrophysiologic stability (Fig. 3B). Based on these findings, 5 µM TH1834 was selected for subsequent experiments as a dose that inhibits TIP60 activity without overtly disrupting basal contractile behavior.

**Figure 3.**
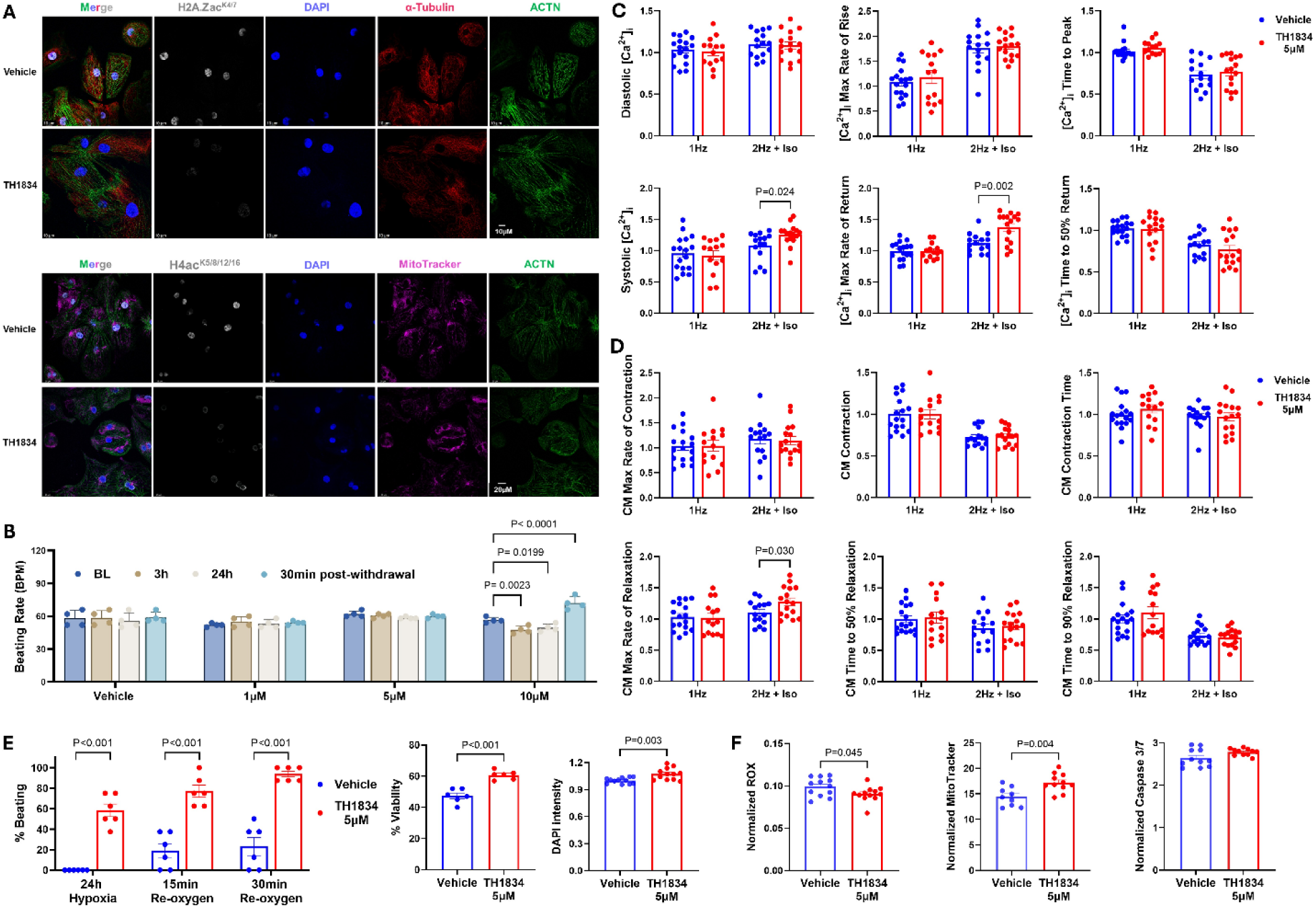
TH1834 inhibits TIP60 acetyltransferase activity and promotes hiPSC-CM survival after H/R injury. **A.** Representative immunofluorescence images of hiPSC-CMs treated overnight with vehicle or 5 µM TH1834 and stained for DAPI, α-tubulin, acetylated histone H4 (H4ac^K5/8/12/16^), acetylated histone H2A.Z (H2A.Zac^K4/7^), sarcomeric α-actinin (ACTN), and MitoTracker. Scale bars are indicated in the images. **B.** Dose-response assessment of TH1834 effects on spontaneous beating behavior under normoxic conditions. hiPSC-CMs were treated with vehicle or increasing concentrations of TH1834, and beating rates were quantified before treatment, at 3 h and 24 h after treatment, and 30 min after drug withdrawal. **C, D.** Effects of 24h TH1834 treatment on single cell intracellular Ca²⁺ ([Ca²⁺]_i_) handling (**C**) and contractile function (**D**). [Ca²⁺]_i_ transient and contraction parameters were quantified from hiPSC-CMs paced at 1 Hz under basal normoxic conditions or paced at 2 Hz during β-adrenergic stimulation with isoproterenol (Iso, 1 µM). **E.** Assessment of post-H/R beating recovery and hiPSC-CM survival following treatment with vehicle or 5 µM TH1834 during H/R. Beating probability was quantified after hypoxia and after 15 min and 30 min reoxygenation. Viable cell number and DNA content (DAPI) were measured after 30min re-oxygenation. **F.** Quantification of reactive oxygen species (ROX), functional mitochondria (MitoTracker), and activated caspase-3/7 after H/R in vehicle- and TH1834-treated hiPSC-CMs. Data are presented as mean ± SEM. Ca²⁺ transient and contraction data were normalized to the averaged values of the 1Hz vehicle group within each experimental batch. Statistical comparisons are indicated in each panel. H/R, 24 h hypoxia at 0.5% O₂ with 3 mM glucose followed by 30min reoxygenation at 21% O₂.

We next examined whether TH1834 alters [Ca²⁺]_i_ handling and contractile function under basal and stress-stimulated conditions. Under normoxic conditions, 24h TH1834 treatment had little effect on kinetics of [Ca²⁺]_i_ or cross-bridge turnover in hiPSC-CMs paced at 1Hz, indicating that TIP60 inhibition does not substantially perturb basal [Ca²⁺]_i_ homeostasis or mechanical performance (Fig. 3C, D). To evaluate β-adrenergic reserve, hiPSC-CMs were stimulated at 2 Hz and challenged with the β-adrenergic receptor agonist isoproterenol (Iso, 1 µM). Under these stress conditions, 24h treatment of TH1834 increased systolic [Ca²⁺]_i_, accelerated [Ca²⁺]_i_ transient decay, and promoted cross-bridge detachment rate, indicative of enhanced β-adrenergic responsiveness and improved relaxation-related kinetics (Fig. 3C, D).

To determine whether TH1834 improves post-ischemic recovery, hiPSC-CMs were treated with 5 µM TH1834 during the optimized H/R protocol. TH1834 increased the probability of spontaneous beating recovery following 24h hypoxia and after 15 min and 30 min reoxygenation, indicating improved functional resilience after ischemic stress (Fig. 3E). Consistent with this functional benefit, TH1834 also improved CM survival, as reflected by increased numbers of viable cells and higher DNA content following H/R injury (Fig. 3E). To further define the protective phenotype, we quantified mitochondrial and stress-related endpoints after H/R. TH1834 treatment during H/R increased the abundance of functional mitochondria and reduced ROS levels, supporting improved mitochondrial integrity and diminished oxidative stress during recovery (Fig. 3F). However, TH1834 had minimal effects on caspase-3/7 activity, suggesting that its protective action is not primarily mediated by strong suppression of apoptotic signaling at this time point.

Together, these data indicate that TH1834 inhibits TIP60 acetyltransferase activity in hiPSC-CMs, preserves basal [Ca²⁺]_i_ and contraction behavior at the selected dose, enhances β-adrenergic functional reserve, and promotes post-ischemic recovery by improving survival, preserving mitochondrial function, and reducing oxidative stress.

### TH1834 preserves Ca^2+^ handling, contractile function, and metabolic capacity in hiPSC-CMs after H/R

To determine whether TH1834 improves functional recovery after ischemic stress, we performed simultaneous real-time single cell measurements of [Ca²⁺]_i_ and contraction in hiPSC-CMs following H/R. TH1834 treatment substantially rescued H/R-induced [Ca²⁺]_i_ defects in CMs stimulated at 1 Hz. TH1834 normalized diastolic [Ca²⁺]_i_, restored [Ca²⁺]_i_ transient decay rate, increased systolic [Ca²⁺]_i_, and shortened the time required for [Ca²⁺]_i_ to return to 50% of baseline (Fig. 4A, B). In addition, TH1834 increased the caffeine-induced SR Ca²⁺ peak, consistent with improved SR Ca²⁺ pump SERCA-mediated Ca²⁺ reuptake and enhanced SR Ca²⁺ load available for release during excitation (Fig. 4B). Together, these findings indicate that TH1834 preserves [Ca²⁺]_i_ homeostasis and improves [Ca²⁺]_i_ cycling efficiency post-H/R. Consistent with Ca²⁺ handling preservation, TH1834 also improved contractile recovery in hiPSC-CMs after H/R. TH1834 restored max systolic contraction, accelerated relaxation, and shortened the time to 90% return to baseline (Fig. 4A, C). These effects indicate rescue of H/R-induced impairment in both cross-bridge turnover and detachment kinetics, supporting improved recovery of excitation-contraction coupling and mechanical performance.

**Figure 4.**
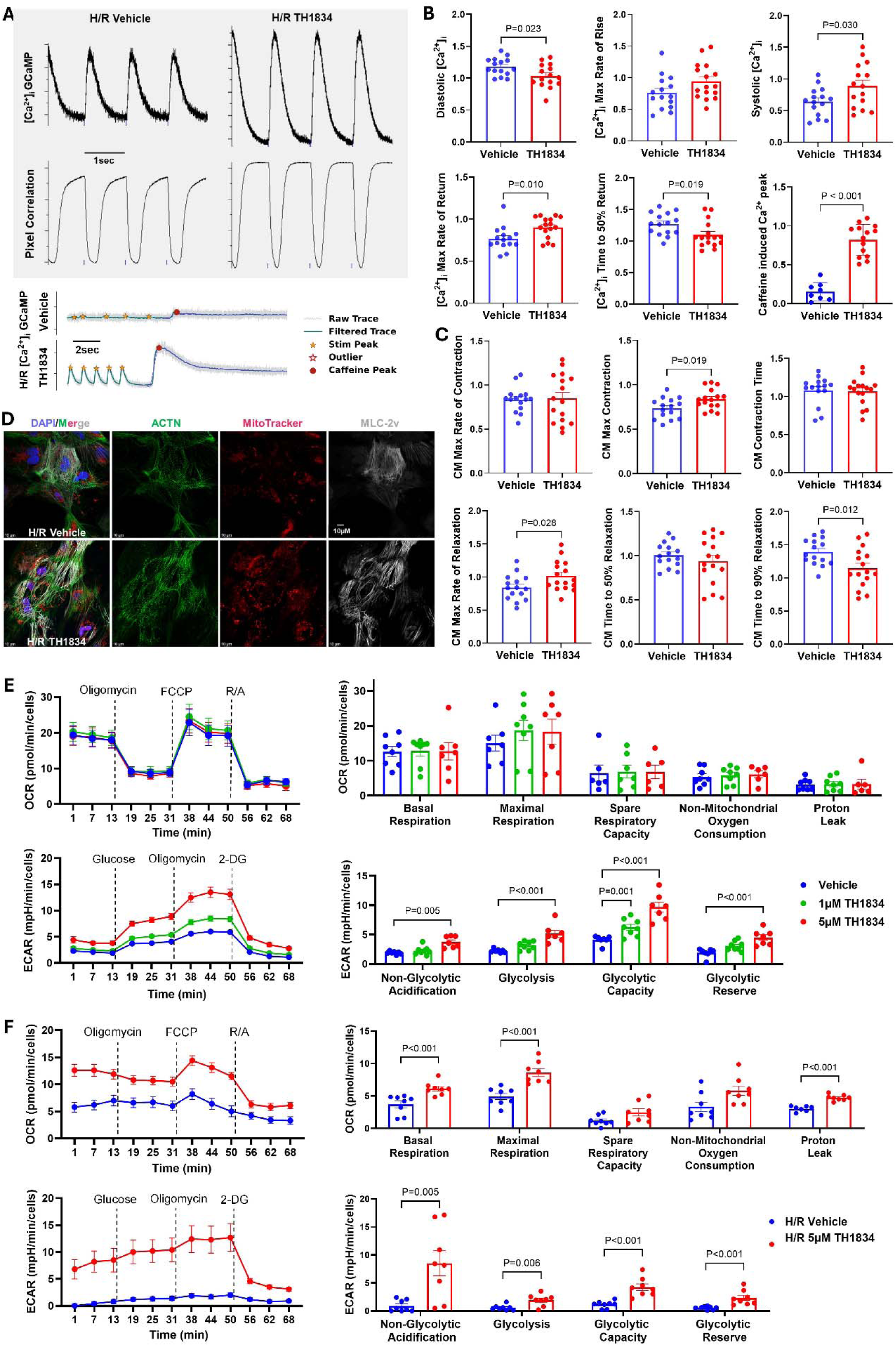
TH1834 preserves Ca^2+^ handling, contractile function, and metabolic capacity in hiPSC-CMs after H/R injury. **A.** Representative simultaneous single-cell recordings of GCaMP-based intracellular Ca²⁺ ([Ca²⁺]_i_) transients and CytoMotion-based contraction paced at 1 Hz (upper) and caffeine-induced sarcoplasmic reticulum (SR) Ca²⁺ release (lower) in vehicle- and TH1834-treated hiPSC-CMs following H/R. **B.** Quantification of [Ca²⁺]_i_ transient parameters following H/R in vehicle- and TH1834-treated hiPSC-CMs paced at 1 Hz and SR Ca²⁺ store. **C.** Quantification of single-cell contraction parameters after H/R in vehicle- and TH1834-treated hiPSC-CMs paced at 1 Hz. **D.** Representative immunofluorescence images of vehicle- and TH1834-treated hiPSC-CMs after H/R stained for DAPI, sarcomeric α-actinin (ACTN), MitoTracker, and ventricular myosin light chain 2 (MLC-2v). Scale bars, 10 µm. **E.** Seahorse Mito Stress and Glycolytic Stress tests of hiPSC-CMs under normoxic conditions following 24h of vehicle, 1 µM TH1834, or 5 µM TH1834 treatment. Oxygen consumption rate (OCR) was measured during sequential injection of oligomycin, FCCP, and R/A, and mitochondria respiration parameters were quantified. Extracellular acidification rate (ECAR) was measured during sequential injection of glucose, oligomycin, and 2-DG, and glycolytic function parameters were quantified. **F.** Seahorse Mito Stress and Glycolytic Stress tests of vehicle- and TH1834-treated hiPSC-CMs after H/R. Data are presented as mean ± SEM. Ca²⁺ transient and contraction data were normalized to the averaged values of the normoxia vehicle group within each experimental batch. Seahorse OCR and ECAR values were normalized to DNA content. Statistical comparisons are indicated in each panel. H/R, 24 h hypoxia at 0.5% O₂ with 3 mM glucose followed by 30min reoxygenation at 21% O₂; TH1834, 5 µM unless otherwise indicated; FCCP, carbonyl cyanide-p-trifluoromethoxyphenylhydrazone; R/A, rotenone/antimycin A; 2-DG, 2-deoxy-D-glucose.

Structural imaging further supported a protective effect of TH1834, as reflected by preserved myofibrillar organization and increased functional mitochondria (Fig. 4D and 3F). These findings suggest that improved Ca²⁺ handling and contractility are accompanied by preservation of sarcomeric and mitochondrial integrity. To further examine the metabolic effects of TH1834, we assessed mitochondrial respiration and cytosolic glycolysis. TH1834 had little effect on mitochondrial respiration under baseline normoxia (Fig. 4E). However, TH1834 increased basal and maximal mitochondrial respiration in hiPSC-CMs following H/R (Fig. 4F), consistent with improved CM viability and increased number of active mitochondria (Fig. 4D, 3E, 3F). Glycolytic stress test revealed that TH1834 increased baseline glycolysis and glycolytic capacity in a dose-dependent manner, with 5 µM TH1834 producing greater effects than 1 µM (Fig. 4E). Post-H/R, elevated glycolytic baseline, capacity, and reserve in TH1834 treated CMs suggests improved anaerobic energy production to support functional recovery (Fig. 4F). These data suggest that although TH1834 does not substantially alter oxidative respiration under normoxia, it enhances glycolytic reserve and metabolic capacity during H/R, supporting energetic recovery following ischemic stress.

Together, these findings demonstrate that 24 h treatment with 5 µM TH1834 preserves Ca²⁺ handling, contractile function, myofibrillar structure, and metabolic adaptability in hiPSC-CMs following H/R injury.

### TH1834 remodels the post-H/R transcriptional state toward metabolic recovery, mitochondrial redox protection, and protein homeostasis

To determine whether the functional protection afforded by TH1834 was accompanied by reversal of H/R-induced transcriptional remodeling, we compared transcriptomes across normoxic, H/R, and TH1834-treated H/R hiPSC-CMs. Among TH1834-responsive genes that moved toward the normoxic state, 442 met stringent rescue criteria (Fig. 5A-C). Strict rescue was strongly directionally asymmetric: 401 genes induced by H/R were reduced by TH1834, whereas only 41 genes suppressed by H/R were significantly increased by TH1834 (Fig. 5D). Thus, TH1834 did not globally normalize the H/R transcriptome but preferentially attenuated H/R-activated injury-associated stress programs.

**Figure 5.**
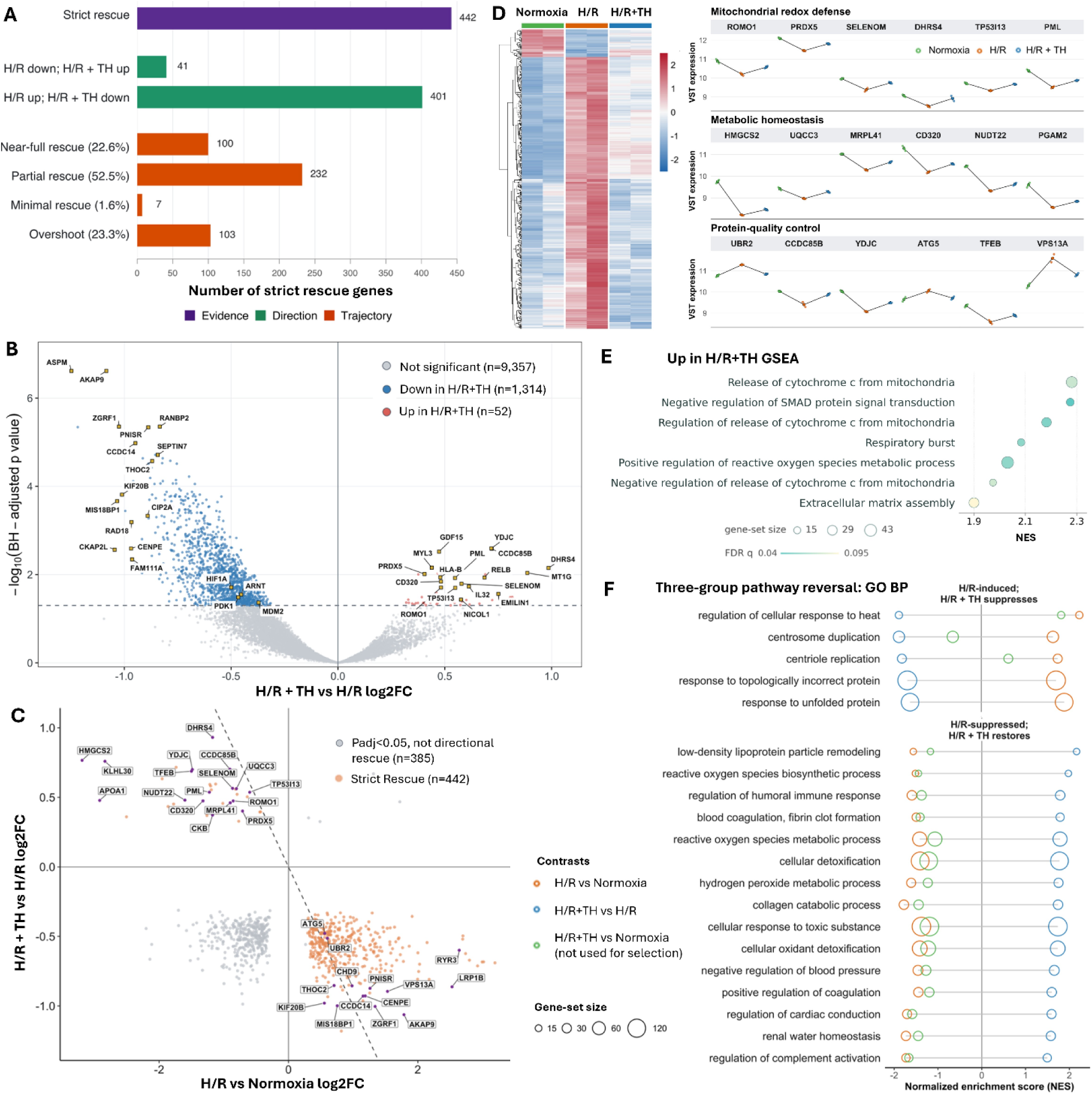
TH1834 remodels the post-H/R transcriptional state toward metabolic recovery, mitochondrial redox protection, and protein homeostasis. **A.** Classification of strict rescue genes. Strict rescue was defined for protein-coding genes with BH-adjusted P<0.05 in both H/R vs normoxia and H/R+TH vs H/R, opposite directions of change across the two contrasts, a concordant VST-expression trajectory, and an H/R+TH group mean closer to normoxia than the H/R mean. Purple indicates the complete strict-rescue set. Green bars separate genes by rescue direction: H/R-down/H/R+TH-up and H/R-up/H/R+TH-down. Orange bars classify the fraction of the H/R-induced expression change reversed by TH1834 as minimal (0–20%), partial (20–80%), near-full (80–105%), or overshoot (>105%). **B.** Volcano plot of differential expression between H/R+TH and H/R. The x-axis shows DESeq2 shrunken log2 fold change and the y-axis shows −log10 BH-adjusted P value. Genes with adjusted P<0.05 are shown in red (increased by TH1834) or blue (decreased); nonsignificant genes are gray. Gold squares denote selected genes highlighted for biological relevance. **C.** Relationship between H/R-induced transcriptional changes and the TH1834 response. Axes show shrunken log2 fold changes for H/R versus normoxia and H/R+TH versus H/R. Orange points indicate strict rescue genes (n=442), whereas gray points indicate genes significant in both contrasts that did not satisfy the strict rescue criteria (n=385). The dashed line (y=−x) denotes equal-magnitude reversal of the H/R effect. **D.** Heatmap of VST-normalized expression for strict rescue genes across normoxia, H/R, and H/R+TH groups, with representative trajectories shown for genes associated with mitochondrial redox defense, metabolic homeostasis, and protein-quality control. Trajectory plots show group-mean VST expression across the three conditions. **E.** Pathway enrichment analysis of the H/R+TH versus H/R transcriptional response. Pre-ranked GSEA of GO Biological Process pathways was performed using genes ranked by the DESeq2 Wald statistic. Gene sets were restricted to 15–500 genes and considered significant at nominal P<0.05 and BH FDR<0.10. Sixteen pathways met these criteria; all seven pathways positively enriched by TH1834 are shown. **F.** GSEA-based identification of H/R-associated pathways reversed by TH1834. Orange denotes H/R vs normoxia, blue denotes H/R+TH vs H/R, and green denotes the residual H/R+TH versus normoxia contrast. Circle position indicates normalized enrichment score (NES), and circle size represents gene-set size. Pathways were retained when H/R and TH1834 produced opposite NES directions and both contrasts met nominal P<0.05 and BH FDR<0.25, with top 20 shown. Green values are shown only to indicate the residual difference from normoxia and were not used for pathway selection.

Direct comparison of H/R+TH1834 with H/R alone further supported this interpretation (Fig. 5B, E). Most significant TH1834-responsive genes were downregulated and involved cell-cycle, DNA damage, and cytoskeletal remodeling, including ASPM, CKAP2L, RAD18, FAM111A, CIP2A, SEPTIN7, and RANBP2. Additional genes with large H/R-induced increases that were reduced by TH1834 included AKAP9, RYR3, LRP1B, CCDC14, ZGRF1, KIF20B, PNISR, MIS18BP1, CENPE, CHD9, and THOC2 (Fig. 5C). Functionally, these genes implicate persistent stress-associated transcriptional, cytoskeletal, Ca²⁺-handling, and injury-remodeling programs after H/R. Their suppression by TH1834 is consistent with attenuation of a maladaptive post-H/R stress state (Fig. 5F). Genes upregulated by TH1834 included MYL3, EMILIN1, RELB, IL32, NICOL1, and CD320, indicating remodeling of cardiac structural integrity, extracellular matrix, and immune-related pathways during recovery.

Several TH1834-responsive genes converged on hypoxia-dependent metabolic adaptation. Compared with H/R alone, TH1834 significantly reduced HIF1A, ARNT, and PDK1 (Fig. 5B). HIF1A and ARNT encode the α and β subunits of HIF-1, respectively, whereas PDK1 is a canonical HIF-1 target that inhibits pyruvate dehydrogenase and restricts mitochondrial entry of glycolysis-derived pyruvate during hypoxia (Kim et al., 2006). Because the transcriptomes were obtained following reoxygenation, attenuation of this HIF-1–PDK1 axis is consistent with reduced persistence of the hypoxic metabolic state during early recovery. TH1834 also restored HMGCS2, UQCC3, MRPL41, CD320, DHRS4, NUDT22, CKB, and APOA1 (Fig. 5C, D), genes linked to mitochondrial metabolism, energy buffering, mitochondrial protein synthesis, and lipid-associated support. Together with improved mitochondrial respiration and glycolytic capacity (Fig. 4), these changes suggest enhanced metabolic adaptability during reoxygenation.

TH1834 also altered genes with functions related to mitochondrial-redox defense and stress adaptation. ROMO1 and PRDX5, which were suppressed by H/R and restored by TH1834 (Fig. 5C, D), are associated with mitochondrial redox regulation and peroxide detoxification (Sabharwal et al., 2013; Xu et al., 2025). Additional H/R-suppressed genes restored by TH1834 included SELENOM, DHRS4, TP53I13, PML, and MT1G, supporting enhanced metal/redox buffering, ER-associated redox homeostasis, metabolic stress adaptation, and p53-linked injury-response signaling during recovery. These changes complement the reduced ROS, increased functional mitochondria, and improved respiratory capacity observed in TH1834-treated hiPSC-CMs after H/R (Fig. 3-4). Genes rescued by TH1834 also implicated protein-quality control and organelle homeostasis, including UBR2, CCDC85B, YDJC, ATG5, TFEB, VPS13A, and KLHL30. (Fig. 5C, D). Thus, TIP60 inhibition appears to reshape mitochondrial, redox, and protein-quality-control signaling during early post-H/R recovery (Fig. 5E, F).

Collectively, the transcriptomic data indicate that TH1834 preferentially suppresses persistent H/R-associated hypoxia, mitotic/replication-stress, DNA-damage, and chromatin-remodeling programs while restoring genes involved in mitochondrial redox protection, metabolic adaptability, protein-quality control, and tissue recovery (Fig. 5A-F). These transcriptional changes parallel the improved mitochondrial integrity, reduced oxidative stress, preserved Ca²⁺ cycling, and enhanced contractile and metabolic recovery observed after TH1834 treatment (Fig. 3-4).

### H/R reproduces hallmarks of ischemia/reperfusion injury in hiPSC-COs

To extend our ischemia/reperfusion model from 2D hiPSC-CMs to a 3D tissue-like context, we next examined the effects of H/R in hiPSC-COs. H/R decreased overall sarcomeric α-actinin (CM) and α-tubulin (CMs and non-CMs) signal intensity (Fig. 6A). We then assessed whether H/R impaired Ca²⁺ handling in hiPSC-COs. H/R significantly reduced systolic [Ca²⁺]_CO_ peak, decreased [Ca²⁺]_CO_ transient rise and decay rates, prolonged time to peak, and delayed return of [Ca²⁺]_CO_ to 50% and 90% of baseline (Fig. 6B). Thus, as in 2D hiPSC-CMs, H/R disrupted Ca²⁺ cycling kinetics in hiPSC-COs, consistent with impaired excitation-contraction coupling after ischemic stress.

**Figure 6.**
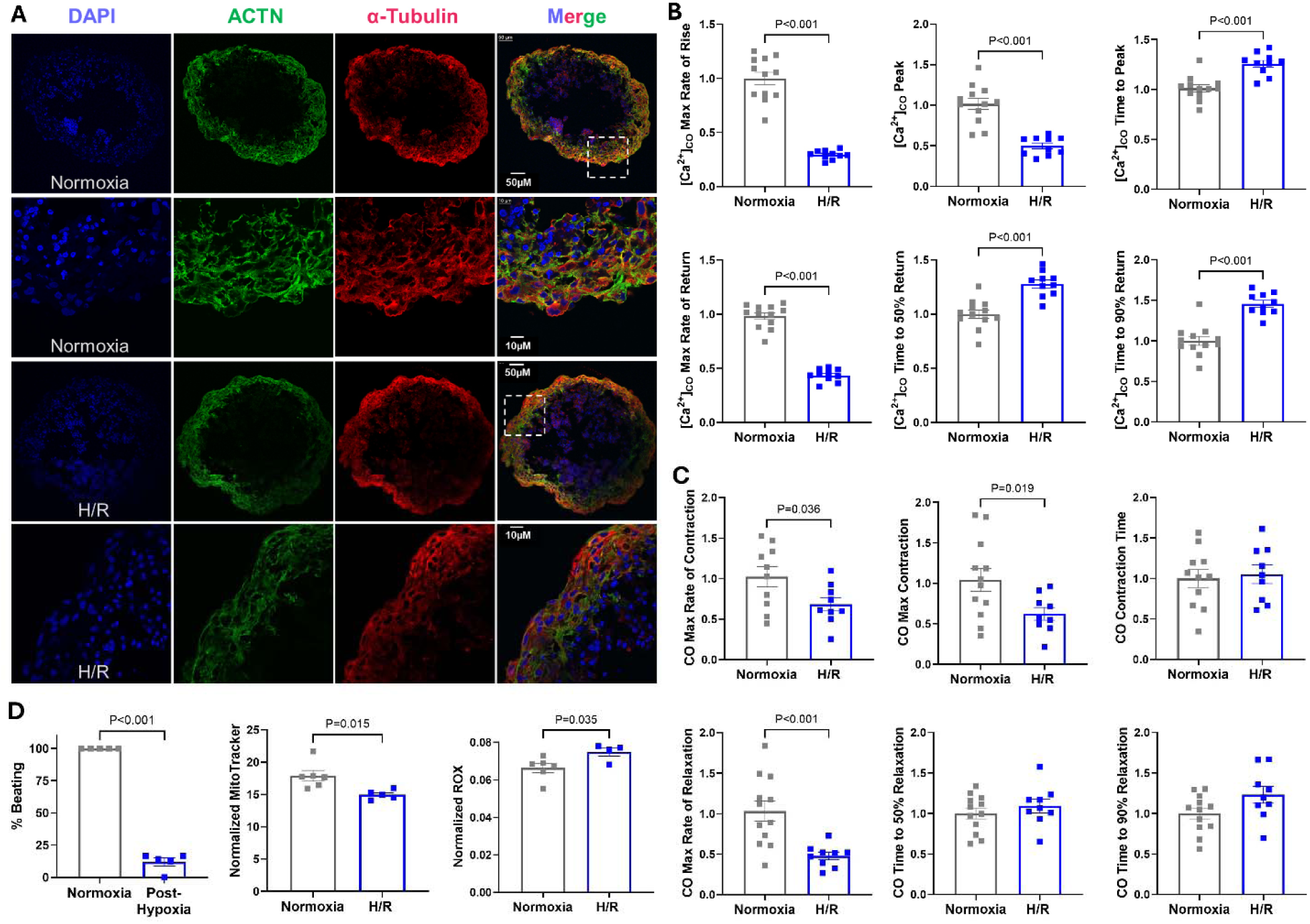
H/R reproduces hallmarks of ischemia/reperfusion injury in hiPSC-COs. **A.** Representative frozen section immunofluorescence images of hiPSC-COs under normoxic conditions or after H/R. Samples were stained for DAPI, sarcomeric α-actinin (ACTN), and α-tubulin to identify CMs and non-CMs. **B.** GCaMP-based Ca²⁺ transient measurements in spontaneously beating hiPSC-COs under normoxic conditions or after H/R. **C.** Brightfield-based contractility measurements in spontaneously beating hiPSC-COs under normoxic conditions or after H/R. **D.** Assessment of spontaneous beating, mitochondria membrane potential, and oxidative stress in hiPSC-COs under normoxic conditions or after H/R. Beating probability was quantified after 24h exposure to hypoxia before re-oxygenation. MitoTracker and CellROX were assessed following H/R. Data are presented as mean ± SEM. Ca²⁺ transient and contraction data were normalized to the averaged values of the normoxia vehicle group within each experimental batch. Statistical comparisons are indicated in each panel. H/R, 24 h hypoxia at 0.5% O₂ with 3 mM glucose followed by 30 min reoxygenation at 21% O₂.

Contractile measurements further showed that H/R markedly depressed mechanical function in hiPSC-COs. Specifically, H/R reduced maximum systolic contraction and velocity and diminished the relaxation rate, demonstrating impaired systolic and diastolic performance in the 3D model (Fig. 6C). These abnormalities indicate that H/R compromises not only Ca²⁺ handling but also integrated tissue-level contractile behavior in hiPSC-COs. Consistent with this injury phenotype, spontaneous beating in hiPSC-COs was arrested following H/R (Fig. 6D). In parallel, H/R reduced the number of functional mitochondria and increased ROS, further supporting substantial metabolic and oxidative injury in the hiPSC-COs (Fig. 6D). Together, these data demonstrate that the H/R model reproduces key hallmarks of ischemia/reperfusion injury in hiPSC-COs, including mitochondrial dysfunction, oxidative stress, impaired Ca²⁺ handling, contractile depression, and loss of spontaneous beating.

### TH1834 preserves Ca^2+^ handling, contractile function, and metabolic recovery in hiPSC-COs after H/R injury

We next asked whether the protective effects of TH1834 extend from 2D hiPSC-CMs to 3D hiPSC-COs subjected to H/R injury. Simultaneous real-time measurements of [Ca²⁺]_CO_ and contraction showed clear functional improvement in TH1834-treated COs compared with vehicle-treated H/R controls (Fig. 7A). TH1834 elevated [Ca²⁺]_CO_ transient departure and return velocities and increased [Ca²⁺]_CO_ peak amplitude (Fig. 7B). These protective effects are in agreement with the enhanced SR Ca²⁺ store and more efficient Ca²⁺ reuptake, observed at the single-cell level in hiPSC-CMs. Consistent with improved [Ca²⁺]_CO_ cycling, TH1834 also rescued H/R-induced contractile dysfunction in hiPSC-COs. TH1834 restored maximum systolic contraction and shortening rate and accelerated relaxation (Fig. 7C). Together, these changes indicate that TH1834 improves both systolic and diastolic performance and reverses the H/R-induced impairment in cross-bridge turnover kinetics.

**Figure 7.**
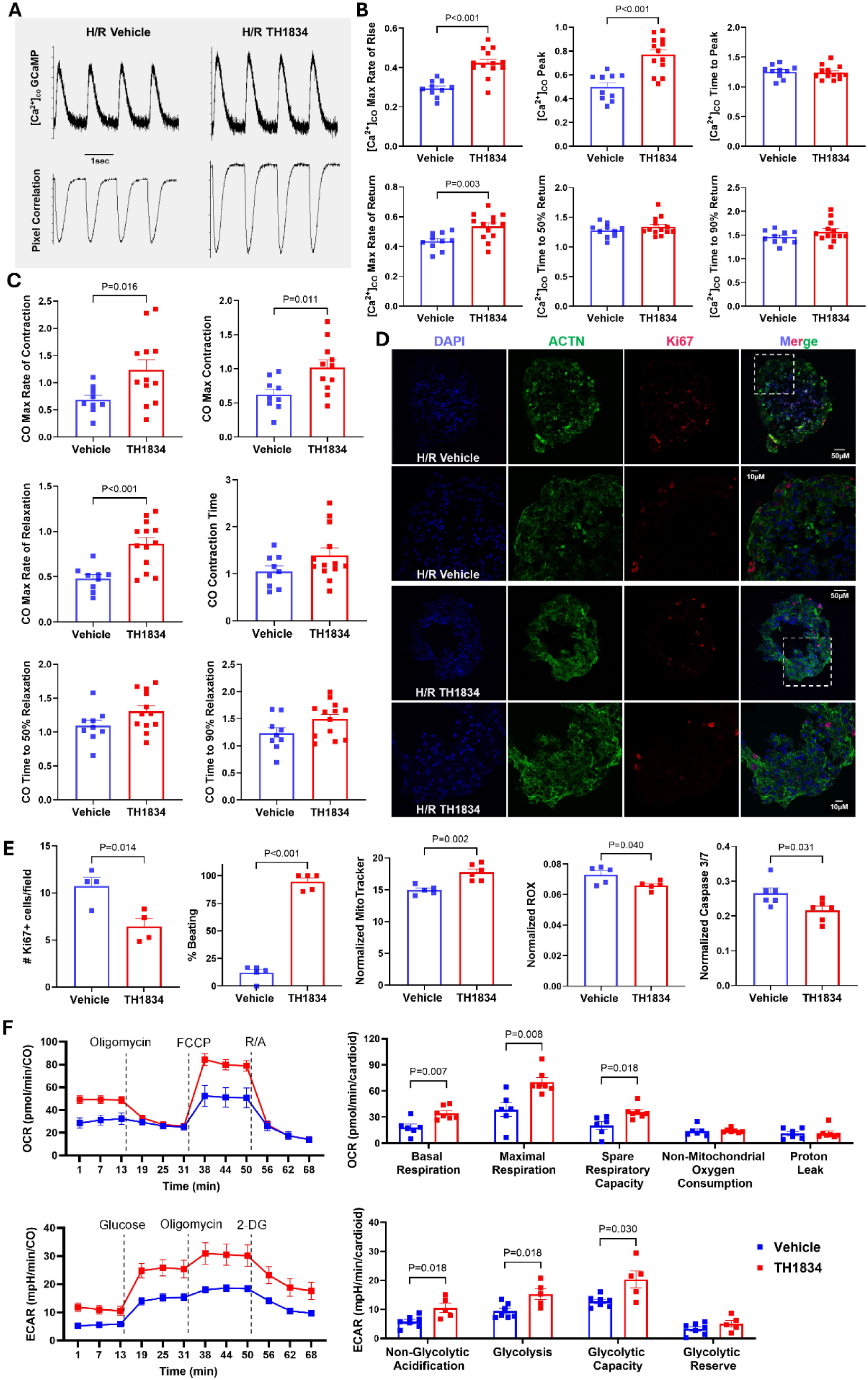
TH1834 preserves Ca^2+^ handling, contractile function, and metabolic recovery in hiPSC-COs after H/R injury. **A.** Representative simultaneous recordings of GCaMP-based Ca²⁺ transients and CytoMotion-based contraction in vehicle- and TH1834-treated hiPSC-COs following H/R. **B & C.** Quantification of [Ca²⁺]_CO_ transient and contractility parameters in vehicle- and TH1834-treated hiPSC-COs after H/R. **D.** Representative immunofluorescence images of frozen hiPSC-CO sections from vehicle- and TH1834-treated groups after H/R stained for DAPI, sarcomeric α-actinin (ACTN), and Ki67. Scale bars are indicated in the images. **E.** Quantification of Ki67-positive cells, spontaneous beating recovery, active mitochondria, reactive oxygen species, and caspase-3/7 activity in vehicle- and TH1834-treated hiPSC-COs after H/R. Spontaneous beating was assessed after 24 h hypoxia, while Ki67, MitoTracker, CellROX, and CellEvent caspase-3/7 were measured after reoxygenation. **F.** Seahorse Mito and Glycolytic Stress Tests on vehicle- and TH1834-treated hiPSC-COs after H/R. Oxygen consumption rate (OCR) was measured during sequential injection of oligomycin, FCCP, and R/A. Extracellular acidification rate (ECAR) was measured during sequential injection of glucose, oligomycin, and 2-DG. Data are presented as mean ± SEM. Ca²⁺ transient and contraction data were normalized to the averaged values of the normoxia vehicle group within each experimental batch. Seahorse OCR and ECAR values were normalized to DNA content. Statistical comparisons are indicated in each panel. H/R, 24h hypoxia at 0.5% O₂ with 3 mM glucose followed by 30min reoxygenation at 21% O₂; TH1834, 5 µM during 24h+30min H/R; FCCP, Carbonyl cyanide-p-trifluoromethoxyphenylhydrazone; R/A, rotenone/antimycin A; 2-DG, 2-deoxy-D-glucose.

TH1834 treatment elevated the overall sarcomeric α-actinin intensity while decreased the number of cells that are positive for Ki67, a marker actively expressed during G1, S, G2, and mitosis phases of the cell cycle (Fig. 7D, E). In parallel, TH1834 promoted recovery of spontaneous beating after 24h exposure to hypoxia, increased the number of functional mitochondria, reduced oxidative stress, and decreased apoptosis following re-oxygenation (Fig. 7E). These findings indicate that TH1834 not only improves functional recovery, but also limits injury-associated oxidative, cell-cycle and cell death responses.

We next examined whether these benefits were accompanied by improved metabolic function. Seahorse extracellular flux analysis showed that TH1834 increased maximal mitochondrial respiration and spare respiratory capacity in hiPSC-COs following H/R (Fig. 7F), consistent with preservation of basal respiration and the increased number of active mitochondria (Fig. 7E, F), with no change in non-mitochondrial oxygen consumption or proton leak. Glycolytic Stress test further demonstrated that TH1834 enhanced cytosolic glycolysis and glycolytic capacity (Fig. 7F), suggesting improved anaerobic energy production during recovery from ischemic stress when oxidative metabolism is compromised. Thus, TH1834 appears to improve energetic resilience in hiPSC-COs by supporting both mitochondrial and glycolytic function.

Together, these data demonstrate that 5 µM TH1834 preserves Ca²⁺ handling, restores contractile performance, promotes metabolic recovery, reduces oxidative stress and cell death, and enhances spontaneous beating recovery in hiPSC-COs following H/R injury.

## Discussion

In this study, we used complementary 2D hiPSC-CMs and 3D hiPSC-COs to determine whether pharmacologic inhibition of TIP60 protects human cardiac cells from H/R injury. TH1834 inhibited TIP60 acetyltransferase activity at a dose that did not disrupt basal Ca²⁺ handling or contraction and promoted recovery across metabolic, Ca²⁺-handling, structural, and mechanical endpoints. Transcriptomic analysis further showed that TH1834 did not globally restore the normoxic transcriptional state but preferentially counter-regulated H/R-induced stress programs while restoring selected mitochondrial redox and protein-quality-control genes. Together with protection in 3D COs, these findings extend our previous mouse studies and support TIP60 inhibition as a strategy for improving post-ischemic functional recovery.

First of all, we established an ischemia/reperfusion model by combining maturation-enhanced hiPSC-CMs with an optimized H/R protocol. Enhancing hiPSC-CM maturation is critical for model development, because it produced sufficient structural organization and metabolic demand to generate a robust H/R injury phenotype. Our optimized maturation medium promoted sarcomere organization, ventricular marker expression, mitochondrial redistribution, and sustained contraction, while transcriptomic profiling supported a maturation-associated ventricular phenotype with residual developmental features. Consistent with prior studies (Feyen et al., 2020; Peters et al., 2022), maturation-enhanced cells were more vulnerable to low-O₂/low-glucose stress than the cells maintained in conventional RPMI/B27 medium, supporting their suitability for modeling ischemic injury. In parallel, our optimized H/R condition increased oxidative stress, disrupted mitochondria and myofibrils, elevated diastolic Ca²⁺, reduced systolic Ca²⁺ transients and SR Ca²⁺ reserve, slowed Ca²⁺ clearance, and impaired contraction and relaxation. These abnormalities are consistent with established mechanisms in which loss of mitochondrial ATP production compromises SERCA- and ion-transport-dependent Ca²⁺ homeostasis, while reoxygenation promotes mitochondrial ROS production, Ca²⁺ overload, and permeability transition (Murphy & Steenbergen, 2008; Garcia-Dorado et al., 2012; Bertero et al., 2024). RNA sequencing provided complementary molecular evidence: H/R induced hypoxia, oxidative/proteotoxic-stress, and glycolytic-adaptation genes while suppressing sarcomeric, Ca²⁺-handling, fatty-acid metabolic, and respiratory programs. Similar transcriptional remodeling has been observed in hypoxic hiPSC-CMs and infarcted human myocardium (Ward et al., 2021; Czosseck et al., 2024; Kuppe et al., 2022). The convergence of transcriptomic and physiological phenotypes supports that H/R disrupts an integrated excitation–contraction–metabolism network.

TH1834 substantially improved this integrated phenotype. Treatment improved Ca²⁺ cycling efficiency, SR Ca²⁺ loading, and contractile dynamics, accompanied by increased functional mitochondria, and reduced ROS. Because ATP production is tightly coupled to Ca²⁺ handling and sarcomere shortening, restoration of mitochondrial energetic capacity provides a plausible basis for improved SERCA-dependent Ca²⁺ reuptake and cross-bridge cycling (Eisner et al., 2017; Bertero et al., 2024). Importantly, TH1834 had little effect on basal Ca²⁺ and mechanical behavior under normoxia but enhanced responses during β-adrenergic stimulation, suggesting that TIP60 inhibition improves functional reserve under increased energetic demand rather than broadly stimulating resting contractility.

A particularly important finding is that TH1834 enhanced metabolic flexibility. During ischemia, glycolysis becomes essential when oxidative phosphorylation is constrained, whereas effective recovery after reoxygenation requires restoration of mitochondrial respiratory capacity (Murphy & Steenbergen, 2008; Zuurbier et al., 2020). Under normoxia condition, TH1834 had little effect on basal mitochondrial respiration but increased glycolytic capacity. Following H/R, treatment enhanced both mitochondrial respiratory capacity and glycolytic function, indicating that TH1834 did not simply shift metabolism toward glycolysis or oxidative phosphorylation; instead, it improved the capacity to utilize both pathways according to oxygen and energetic availability. Transcriptomic analysis provided further mechanistic support for the improved mitochondrial respiration observed after H/R. TH1834 preferentially counter-regulated H/R-induced transcriptional changes, with 401 of 442 strict rescue genes following an H/R-up/TH1834-down pattern. Among treatment-responsive genes, HIF1A, ARNT, and PDK1 were reduced, suggesting reduced persistence of hypoxia-associated metabolic signaling during reoxygenation (Kim et al., 2006). Additionally, the increase in HMGCS2, UQCC3, CKB, MRPL41, CD320, DHRS4, NUDT22, and APOA1 further supports metabolic remodeling involving mitochondrial substrate utilization, energy buffering, and mitochondrial translation. Together with the preservation of glycolytic capacity, these findings suggest that TH1834 promotes metabolic recovery without forcing a fixed metabolic state, thereby supporting both anaerobic and oxidative energy production as oxygen availability changes.

TH1834 also restored ROMO1 and PRDX5, providing a potential molecular link to improved mitochondrial redox resilience. ROMO1 has recently been shown to protect the mitochondrial cysteinome from oxidation and to support mitochondrial energetics, Ca²⁺ handling, and resistance to permeability transition, including protection of adult CMs from oxidant injury (Xu et al., 2025). PRDX5 similarly limits mitochondrial oxidant signaling during hypoxia, while MT1G and SELENOM suggest additional metal/redox buffering and ER-associated redox homeostasis (Sabharwal et al., 2013). TH1834 also increased GDF15, which protects myocardium against experimental I/R injury (Kempf et al., 2006). These transcriptional changes are consistent with the observed reduction in ROS and preservation of functional mitochondria following TH1834 treatment, suggesting that TIP60 inhibition may promote recovery in part by strengthening mitochondrial and redox defense mechanisms.

One interesting finding is the remodeling of protein-quality-control and stress-response pathways. TH1834 partially restored TFEB, normalized ATG5, and returned the E3 ubiquitin ligase UBR2 toward its normoxic level. TFEB-dependent lysosomal/autophagic function has been implicated in myocardial I/R protection, whereas UBR2 regulates ubiquitin-dependent protein turnover (Gu et al., 2020; Gao et al., 2022). These observations suggest that TH1834 may improve handling of damaged proteins and organelles during recovery, although transcript abundance alone cannot establish autophagic flux or proteasomal activity. Cell-cycle, mitotic, DNA-damage, and chromatin-remodeling pathways were also broadly suppressed by TH1834. This is consistent with reduced Ki67-positive cells in H/R-treated COs and suggests attenuation of injury-associated cell-cycle/stress signaling.—Additional changes suggest remodeling of protein-quality-control, extracellular matrix, contractile, inflammatory, and immune-associated signaling during post-H/R recovery.

The protection observed in 3D hiPSC-COs strengthens the relevance of the 2D findings. COs provide multicellular organization and tissue-level mechanical behavior not captured by CM monolayers and have increasingly been used to model human myocardial injury (Richards et al., 2020; Zhang et al., 2025). H/R impaired mitochondrial function, Ca²⁺ cycling, contractility, and spontaneous beating in the COs, whereas TH1834 improved Ca²⁺ transient kinetics, systolic and diastolic performance, beating recovery, mitochondrial function, and survival. Importantly, TH1834 also enhanced mitochondrial respiratory reserve and glycolytic capacity, reproducing the metabolic flexibility observed in 2D hiPSC-CMs. Thus, the protective phenotype was retained in a more complex human cardiac microtissue environment.

Several limitations should be considered. Maturation-enhanced hiPSC-CMs remain less mature than adult ventricular CMs, and the H/R model cannot reproduce vascular flow, immune-cell interactions, mechanical loading, acidosis, or regional nutrient gradients present in vivo. The RNA-seq experiments also included a limited number of biological replicates and captured an early reoxygenation time point; additional time points will be needed to distinguish transient stress resolution from sustained transcriptional remodeling. Although TH1834 reduced TIP60-dependent histone acetylation, the direct substrates responsible for protection remain unknown. Defining relevant non-histone TIP60 substrates will therefore be important for understanding how transcriptional and post-translational mechanisms converge to produce cardioprotection. Likewise, the multicellular composition of COs complicates assignment of cell-cycle and other responses to individual cell populations. Future studies incorporating additional hiPSC backgrounds, cell-type-resolved analyses, longer recovery periods, metabolic-flux measurements, and acetylomic profiling will help define the mechanism and translational robustness of TIP60 inhibition.

In summary, TH1834-mediated TIP60 inhibition promotes recovery from H/R injury in both 2D human hiPSC-CMs and 3D COs. This protection is characterized by improved metabolic flexibility and mitochondrial-redox resilience, preservation of Ca²⁺ handling and SR Ca²⁺ reserve, maintenance of myofibrillar integrity, and recovery of contraction and relaxation function. Rather than globally normalizing the H/R-induced transcriptomic response, TH1834 selectively attenuates persistent stress programs while restoring mitochondrial-redox defense and protein quality control pathways. The resulting metabolic flexibility may provide the energetic support required to preserve Ca²⁺ reuptake, cross-bridge cycling, and mechanical recovery during the transition from hypoxia to reoxygenation. Together, these findings support TIP60 as a therapeutic target for ischemic heart disease and identify metabolic flexibility as a central feature of TH1834-mediated post-ischemic recovery.

## Supporting information

Supplemental Video 1. RPMI ctrl day 3

Supplemental Video 2. MM day 3

Supplemental Video 3. RPMI ctrl day 6

Supplemental Video 4. MM day 6

Supplemental Video 5. RPMI ctrl day 9

Supplemental Video 6. MM day 9

Supplemental Video 7. RPMI ctrl day 12

Supplemental Video 8. MM day 12

