## Supplemental Table 1 for "TH1834-Mediated TIP60 Inhibition Protects Against Ischemia/Reperfusion Damage in Human iPSC-Derived Cardiomyocytes and Cardioids"

**Supplementary Table S1**. Transcriptional markers of normoxic maturation-enhanced hiPSC-CMs

| **Functional module** | **Gene** | **Protein** | **Normoxia TPM** | **Normoxia FPKM** |
| --- | --- | --- | --- | --- |
| **Sarcomere / contractile** | ACTC1 | Cardiac α-actin | 10,472.77 | 2,321.78 |
|  | TNNC1 | Cardiac troponin C | 2,048.54 | 456.44 |
|  | TNNT2 | Cardiac troponin T | 41.62 | 9.21 |
|  | TNNI3 | Adult cardiac troponin I | 45.79 | 10.14 |
|  | TNNI1 | Fetal/slow skeletal troponin I | 563.19 | 124.99 |
|  | MYH7 | β-myosin heavy chain | 830.25 | 184.46 |
|  | MYH6 | α-myosin heavy chain | 1,230.09 | 272.20 |
|  | MYL2 | Ventricular MLC-2v | 148.47 | 33.10 |
|  | MYL7 | MLC-2a | 5,139.67 | 1,140.54 |
|  | MYL3 | Myosin essential light chain | 830.61 | 184.56 |
|  | MYBPC3 | Cardiac myosin-binding protein C | 699.60 | 154.95 |
|  | ACTN2 | Sarcomeric α-actinin | 125.37 | 27.72 |
|  | TTN | Titin | 82.48 | 18.22 |
|  | MYOM1 | Myomesin-1 | 129.73 | 28.67 |
|  | MYOM2 | Myomesin-2 | 0.73 | 0.16 |
|  | MYOM3 | Myomesin-3 | 0.40 | 0.089 |
| **Ventricular identity** | IRX4 | Ventricular transcription factor | 24.04 | 5.34 |
|  | HEY2 | Ventricular/compact-myocardium TF | 17.04 | 3.78 |
|  | TBX5 | Cardiac chamber TF | 12.42 | 2.76 |
| **SR Ca²⁺ handling / E–C coupling** | ATP2A2 | SERCA2 | 260.81 | 57.64 |
|  | JPH2 | Junctophilin-2 | 56.07 | 12.44 |
|  | SLC8A1 | NCX1 | 74.87 | 16.52 |
|  | CACNA1C | Cav1.2 α1C | 16.80 | 3.72 |
|  | CACNB2 | L-type Ca²⁺ channel β2 | 12.83 | 2.83 |
|  | TRDN | Triadin | 7.96 | 1.76 |
|  | RYR2 | Ryanodine receptor 2 | 35.83 | 7.92 |
|  | PLN | Phospholamban | 203.45 | 45.04 |
|  | CASQ2 | Calsequestrin-2 | 0.61 | 0.13 |
|  | S100A1 | Cardiac Ca²⁺ regulatory protein | 1.08 | 0.24 |
| **Electrical coupling / ion channels** | GJA1 | Connexin-43 | 321.97 | 71.52 |
|  | SCN5A | Nav1.5 | 25.18 | 5.58 |
|  | KCNJ2 | Kir2.1 / I_K1_ | 6.07 | 1.35 |
|  | KCNH2 | hERG / I_Kr_ | 56.14 | 12.45 |
|  | KCNQ1 | Kv7.1 / I_Ks_ | 8.00 | 1.78 |
|  | KCND3 | Kv4.3 / I_to_ | 0.54 | 0.12 |
|  | HCN4 | Pacemaker current | 100.12 | 22.25 |
| **Energy transfer / mitochondrial maturation** | CKM | Muscle creatine kinase | 406.17 | 90.15 |
|  | CKMT2 | Mitochondrial creatine kinase | 16.49 | 3.66 |
|  | COX6A2 | Heart/muscle cytochrome-c oxidase subunit | 22.54 | 5.01 |
|  | FABP3 | Cardiac fatty-acid binding protein | 84.34 | 18.70 |
|  | PPARA | PPARα | 15.19 | 3.36 |
|  | ESRRA | ERRα | 60.83 | 13.48 |
|  | NRF1 | Nuclear respiratory factor 1 | 13.76 | 3.06 |
|  | TFAM | Mitochondrial transcription factor A | 27.93 | 6.20 |
|  | CPT1B | Mitochondrial FA import | 0.40 | 0.087 |
|  | ACADM | Medium-chain acyl-CoA dehydrogenase | 11.17 | 2.48 |
|  | HADHA | Mitochondrial trifunctional protein α | 146.75 | 32.55 |
|  | HADHB | Mitochondrial trifunctional protein β | 74.64 | 16.55 |
|  | PPARGC1A | PGC-1α | 20.04 | 4.43 |
|  | CD36 | Fatty-acid transporter | 0.41 | 0.091 |
|  | ACADVL | Very-long-chain acyl-CoA dehydrogenase | 247.99 | 55.00 |
| **OXPHOS / mitochondrial capacity** | NDUFS1 | Complex I | 61.47 | 13.63 |
|  | UQCRC1 | Complex III | 239.90 | 53.23 |
|  | ATP5F1A | ATP synthase α | 309.44 | 68.66 |
|  | SLC25A4 | ANT1 | 230.62 | 51.07 |
| **Glycolytic state** | HK2 | Hexokinase-2 | 152.56 | 33.78 |
|  | LDHA | Lactate dehydrogenase A | 949.39 | 210.29 |
